# Patient-derived functional immuno-oncology platform identifies responders to ATR inhibitor and immunotherapy combinations in ovarian cancer

**DOI:** 10.1101/2024.02.15.579904

**Authors:** Ashwini S Nagaraj, Matilda Salko, Aditi Sirskikar, Erdogan Pekcan Erkan, Elina A Pietilä, Iga Niemiec, Jie Bao, Giovanni Marchi, Angéla Szabó, Kirsten Nowlan, Sanna Pikkusaari, Anna Kanerva, Johanna Tapper, Riitta Koivisto-Korander, Liisa Kauppi, Sampsa Hautaniemi, Anna Vähärautio, Jing Tang, Ulla-Maija Haltia, Eliisa Kekäläinen, Anni Virtanen, Tuula Salo, Anniina Färkkilä

## Abstract

Responses to single agent immunotherapies have remained modest in high-grade serous ovarian cancer (HGSC), suggesting the need for combination treatments. Identifying clinically effective immunotherapy combinations (IC) requires pre-clinical testing using models representing the patient-specific immune microenvironment. Here, we established a functional immuno-oncology platform for high-throughput and functional testing of IC using HGSC patient-derived immunocompetent cultures (iPDCs) established on patient-derived omentum gel matrix. We employed genomic and single-cell analysis to assess the intricate and functional characteristics of the iPDCs combined with tumor and immune cell-specific cytotoxic responses. Corroborating the clinical response to Poly (ADP-ribose) polymerase inhibitors (PARPi), iPDCs showed homologous recombination deficiency (HRD) - specific response to PARPi. Importantly, drug responses from iPDCs of chemotherapy and PARPi refractory patients corresponded with patient outcomes and aligned with distinct pathway activities from single-cell RNA sequencing analysis. Furthermore, iPDCs from HRD tumors showed response to anti-PD1 antibody as measured by decrease in tumor cells combined with augmented T cell activation. High-throughput drug testing followed by single cell-imaging from iPDCs revealed patient-specific responses to combination of ataxia telangiectasia and Rad3-related inhibitor (ATRi) with DNA damaging agents or immunotherapies. Integration of cytotoxic responses with immune cell states uncovered patient-specific immune activation with the combination of ATRi and a novel immunotherapy targeting Autotaxin (ATX), and this response was significantly associated with a tumor-cell replication stress biomarker in single-cell analysis of tCycIF highly multiplexed imaging. In conclusion, iPDCs provide a platform for high-throughput screening and functional testing of immuno-oncology agents for precision oncology in HGSC.

## INTRODUCTION

High-grade serous ovarian carcinoma is the most common subtype of ovarian cancer, often diagnosed at an advanced stage with 10-year survival of 15% (*1*). Nearly 80% of the HGSC metastasize to omentum, a large fat tissue covering stomach and intestine, which provides metabolic and inflammatory microenvironment for cancer and immune cells (*2*). The standard of care for HGSC includes primary debulking surgery followed by platinum-based chemotherapy. Despite the successful response to initial treatment, recurrent disease often develops within the first three years. Poly (ADP-ribose) polymerase inhibitors (PARPi) have improved patient’s outcome when used as a first-line or maintenance therapy, particularly in patients with homologous recombination deficiency (HRD), and or sensitivity to platinum-based chemotherapy (*3*). However, 35-45% of the patients discontinue the treatment due to innate or acquired resistance to PARPi (*4, 5*), suggesting that novel combination strategies are required to overcome resistance to PARPi.

Single agent immune checkpoint inhibitors (ICIs) have shown limited efficacy in HGSC with overall response rate of 10% (*6*). The lack of success with ICIs in HGSC is likely due to a variety of immunosuppressive cells and mechanisms in the tumor microenvironment (TME) (*7*). Targeting immunosuppressive mechanisms operated by myeloid cells is one of the strategies to improve the efficacy of ICIs. Towards this, studies from mouse models show the importance of targeting Autotaxin-*Lysophosphatidic acid (LPA*) pathway or PKR-like endoplasmic-reticulum (ER) kinase (PERK) to enhance CD8+ T cell response via activating interferon signaling in dendritic cells (DCs) or by depleting myeloid-derived suppressor cells (MDSC), respectively (*8, 9*).

To improve the therapeutic benefit of ICIs, their combination with chemotherapy, PARPi, or anti-angiogenic agents are under clinical investigation (*10, 11*). Interestingly, results from the TOPACIO Phase I/II clinical trial showed that mutational signature 3 as a surrogate of HRD status predicts sensitivity to combination of PARPi, niraparib and anti-PD1 antibody, pembrolizumab (*12*). In addition, *BRCA 1/2* defective HGSC has been shown to exhibit increased immunosurveillance with more neoantigens, tumor infiltrating lymphocytes, programmed cell death (PD-1) and its ligand (PD-L1) compared to HRP tumors (*13–15*). These findings highlight the investigation of novel immunotherapy combinations for patients with HGSC. Various agents inhibiting DNA damage response (DDRi) including cell cycle checkpoint inhibitors targeting ATR or WEE1 kinases are being clinically explored (*16*). While preclinical studies implicate benefit of combining of DDRi and ICIs (*17*), investigation of novel combinations of ATRi with other DNA damaging agents or immunotherapies is required to guide personalized immuno-oncology in ovarian cancer. However, prior to clinical investigation of novel immunotherapies, the efficacy of these agents needs to be validated in immunocompetent models representing patient tumor-specific immune microenvironment.

A variety of patient-derived *ex vivo* models have been developed to model tumor immune microenvironment, and to evaluate response to immunotherapies (*18–20*). All these studies have utilized matrigel or collagen-based matrices to culture the cells or tissue fragments. However, the source of these matrices being commonly of mouse or rat origin, do not represent HGSC patient TME. In addition, previous studies using patient-derived models have primarily focused on the effect of ICIs on functional responses from immune cells but not the cytotoxic response from tumor cells. Furthermore, these models are not suitable for high-throughput combinatorial drug testing. To address this, we established a functional precision immuno-oncology platform using novel HGSC patient-derived immunocompetent cultures (iPDCs) on a patient-derived omentumgel (OmGel) matrix produced from tumor-free omentum tissue (*21*). We validated that iPDCs preserve tumor and immune cell features of the native tumor and exhibit previously shown HRD tumor-specific response to PARPi. Further, we provide novel insight into HRD-specific sensitivity to anti-PD1 antibody treatment. In addition, we show that iPDCs can reveal drug sensitivity and resistance and recapitulate clinical responses to PARPi, or chemotherapy. Furthermore, by coupling single cell image-based tumor and immune cell-specific drug responses with immune cell functional responses we show that iPDCs reveal patient-specific vulnerabilities to immuno-oncology agents and combination therapies in HGSC.

## RESULTS

### Patient-derived novel OmGel matrix significantly improves the growth of HGSC patient-derived cultures that are applicable for drug testing

In this study, we have processed 42 HGSC tumors from 39 patients and established cultures from 38 tumors (**Table. S, 1**). We first started with optimizing tumor tissue dissociation, as the enzymatic dissociation is a necessary step in setting up *ex vivo* cultures and can influence on the successful establishment of patient tumor-derived cultures. We compared two commonly used tissue dissociating enzymes: Collagenase/Hyaluronidase (C&H), and dispase for the final yield, viability, abundance of tumor and immune cells, and expression of stress response genes (**Fig. S1, A**). Adjacent tissue regions from each tumor were dissociated with C&H or dispase. Our results showed no significant difference in the final yield or calcein positive live cells between C&H and dispase-dissociation (**Fig. S1, B-D**). Similarly, the abundance of CK7+ tumor cells or CD45+ immune cells were not affected by the nature of enzymes (**Fig. S1, E and F**). However, the expressions of stress response genes *ATF3*, *FOS*, and *HSPA1* were increased in C&H dissociated cells compared to dispase (**Fig. S1G**). The upregulation of stress response genes has been linked to chemoresistance (*22, 23*) and therefore can distort drug response analysis from cancer cells. Hence, we continued with dispase-dissociation for establishing iPDCs (**Fig. 1, A**). HGSC primary or omental metastatic tumor-dissociated cells were cultured in basement membrane extract (BME) or OmGel for up to one week. Compared to cells cultured in BME, OmGel cultured cells showed significantly improved growth as measured by the number of PDCs with size >3000 µm^2^ (**Fig. 1, B, C, and S1. H**). Importantly, OmGel cultured PDCs maintained source tumor morphology and expression of tumor-specific marker PAX8 (**Fig. 1, D and E**). In addition, whole exome sequencing (WES) analysis of tumor and matched PDCs (n=10) showed strong correlation (r=0.86) for the variant allelic frequencies (VAF), and a high median concordance (95.8%) for tumor-specific mutations between tumor and PDC pairs (**Fig. 1, F-G**, **Table. S, 2**).

**Fig. 1.**
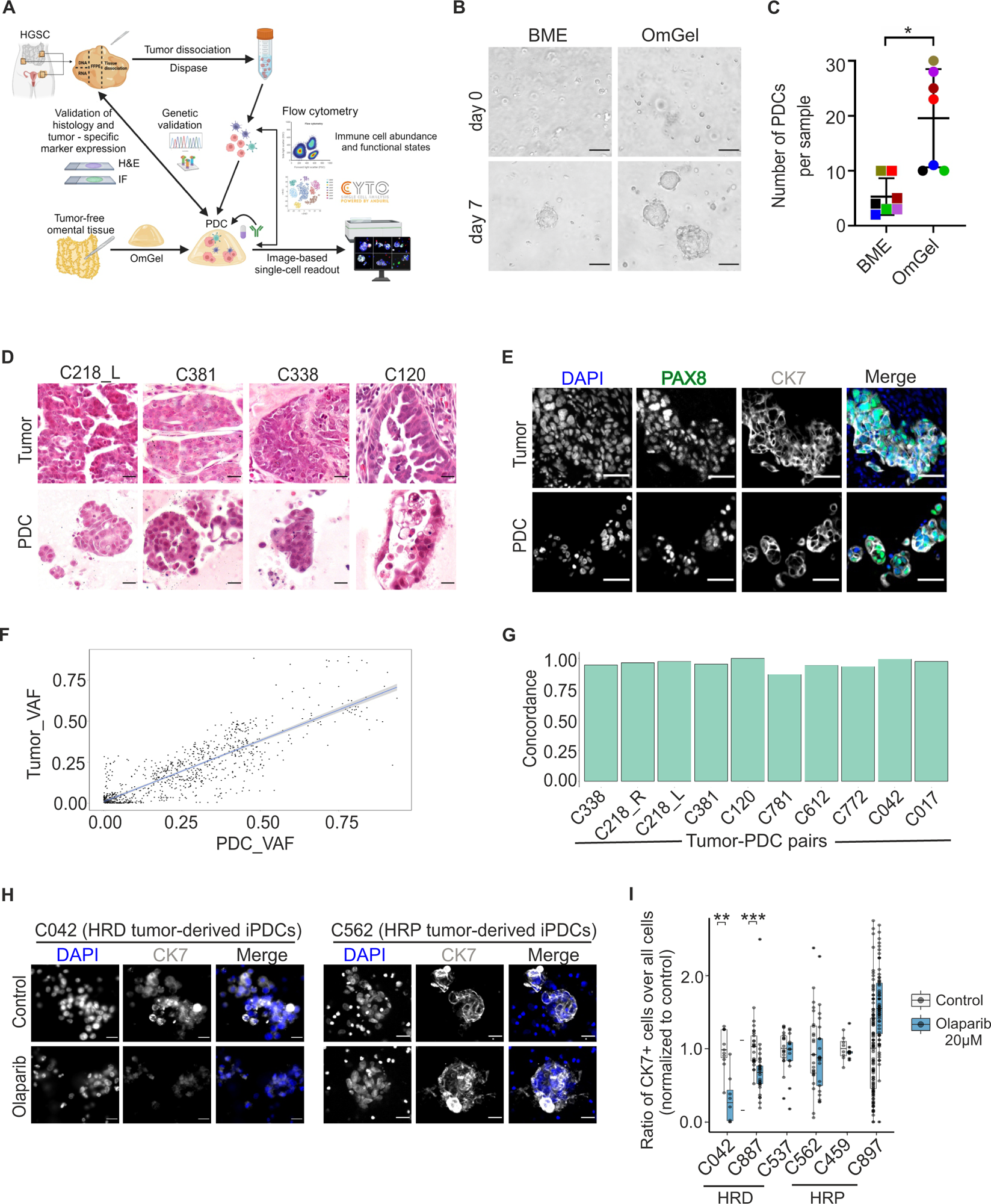
Patient-derived novel OmGel matrix enables significantly improved growth of HGSC patient-derived cells that retain source tumor-specific features and mimic clinical response to PARPi. **A).** Schematics showing the establishment and validation of iPDCs cultivated in OmGel, followed by drug testing on iPDCs and drug response analysis using image-based single cell readouts. **B).** Representative brightfield images showing HGSC patient-derived cells cultured in basement membrane matrix (BME) or OmGel. Scale bar 50µM. **C).** Quantification of number of PDCs cultured in BME or OmGel for 4-10 days, each color represents an individual sample. **D).** Representative H&E images of the tumor tissue and matched PDCs cultured in OmGel for 4-6 days. Scale bar 100µM. **E).** Representative IF images of tumor tissue and matched PDCs cultured in OmGel and stained with indicated antibodies. Scale bar 50µM. **F).** Scatter plot depicting the correlation of variant allele frequencies (VAF) between tumor and PDCs. **G).** Bar plot representing the concordance between tumor tissue and matched PDCs for tissue-specific mutations. **H).** Representative IF images of control or olaparib-treated iPDCs stained with CK7 antibody. Scale bar 50µM. **I).** Box plot showing the ratio of CK7+ cells over all cells normalized to control. Analysis was performed 4-6 days post olaparib treatment. Each dot represents an individual image. Error bars represent mean ± standard deviation (SD). Wilcoxon signed rank test * p< 0.05, ** p< 0.01 *** p<0.001.

Next, we examined whether OmGel cultured PDCs could be utilized for drug response analysis. Clinically PARP inhibitor maintenance therapy is effective particularly in HGSC patients with HRD and or *BRCA* mutation (*24*). However, nearly 30% of the patients with a germline *BRCA* mutation relapse within the first 2 years on a PARPi maintenance therapy (*25*). We tested if the PDCs established from treatment naive patient-derived tumors exhibit HRD genotype-specific responses to PARPi. All the tumor samples used in this study were subjected to the genomic ovaHRDscar test to evaluate homologs recombination (HR) status (**Table. S, 3**) (*26*). We treated the treatment naive HRD or HRP tumor derived PDCs (n=3 each) with olaparib and evaluated the treatment responses as a decrease in the abundance of CK7+ tumor cells compared to the control. While 2/3 HRD-PDCs (C042 and C887) showed a significant decrease in the abundance of CK7+ cells, none of the HRP tumor-derived PDCs showed significant responses upon olaparib treatment (**Fig. 1, H and I**). Interestingly, the HRD-PDCs (C537) that did not respond to olaparib had a HRD score of 56 close to the cut-off value (55), while the responding PDCs had higher HRD scores (C042: 72, C887:65), thus potentially reflecting the relative abundance of HR deficient cells in PDCs from C537. One of the patients (C042), whose PDCs responded to PARPi, also exhibited clinical benefit without evidence for disease progression after 10 months of PARPi maintenance therapy. For the remaining patients, clinical PARPi responses were unavailable. In summary, these results show that novel OmGel matrix enable successful generation of HGSC PDCs that are amenable for drug response analysis and that the PDC drug responses relate to the genotype-predicted responses to PARPi therapy.

### Patient-derived immunocompetent cultures maintain native immune cells and show HRD-specific responses to anti-PD1 antibody treatment

To evaluate if the nature of the enzyme or culturing matrix influence on the recovery of immune cell types, we performed multi-parameter flow cytometry on C&H or dispase-dissociated cells or cells cultured with BME or OmGel. The populations negative for dead cell marker and epithelial cells (DCM^-^ EpCAM^-^), were gated in FlowJO and further analyzed using CYTO (*27*) (**Fig. 2, A and Fig. S2, A**). The DCM^-^ EpCAM^-^ data from all the samples comparing enzymes or matrix conditions were clustered into 25 different clusters using FlowSOM, and annotated as T cell, myeloid, or other clusters. The data from these clusters were further analyzed using CYTO and annotated as different immune cell types (**Fig. S2, A**). Comparison of the immune cell proportions between C&H vs dispase-dissociation or between BME vs OmGel cultures did not show statistically significant difference for any of the cell types tested using paired wilcoxon signed rank test (**Fig. S2, B and C**).

**Fig. 2.**
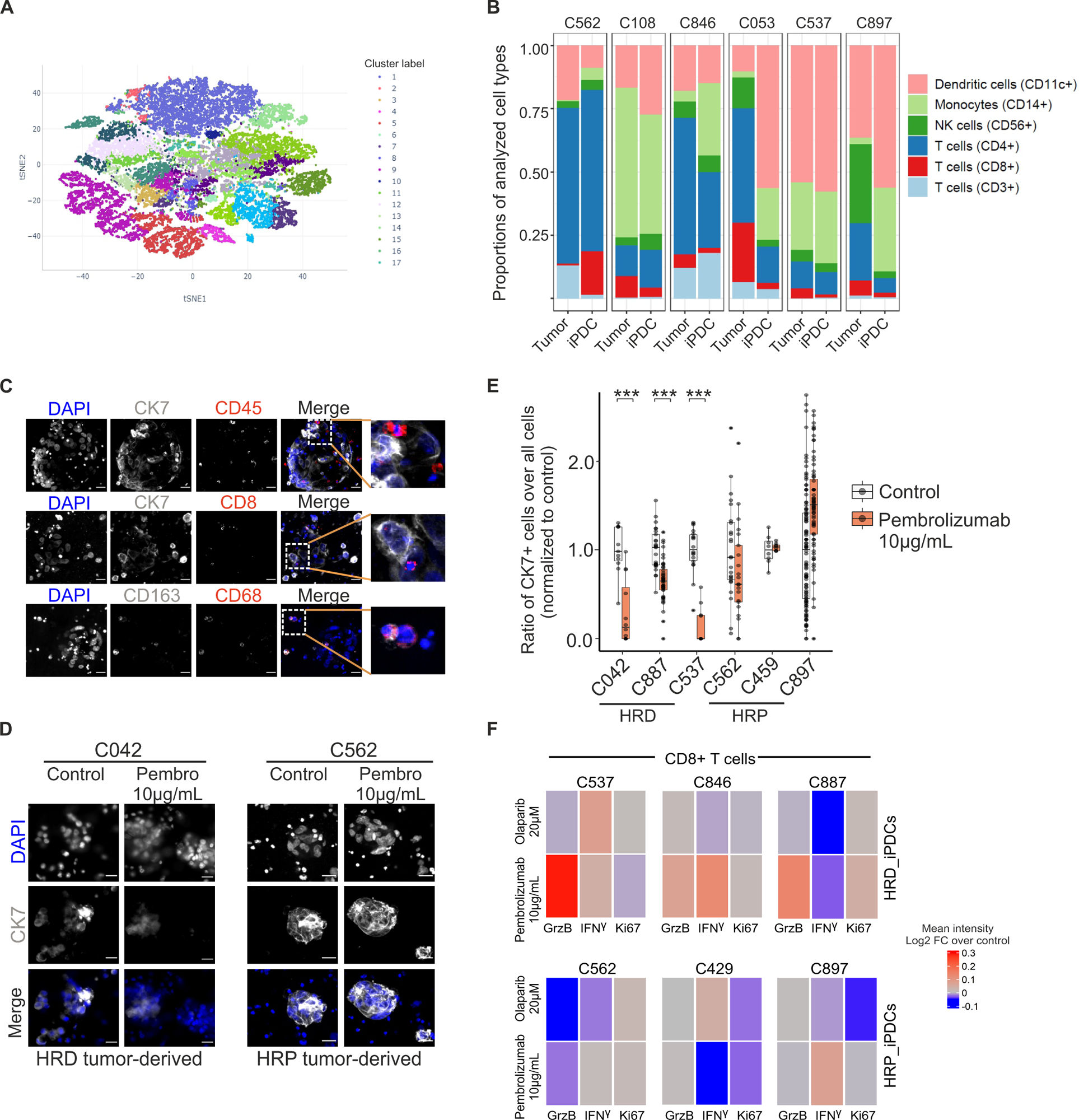
HGSC patient-derived OmGel cultures retain source tumor-derived immune cells and exhibit HRD genotype-specific response to anti-PD1antibody treatment. **A).** t-SNE plot depicting the clusters analyzed from EpCAM^-^ cells of tumor and matched iPDCs (n=6), cultured for 4-6 days. **B).** Bar plot showing the proportions of indicated immune cells in tumor and matched iPDCs, annotated from the clusters shown in (A) **C).** Representative IF images of day 4 iPDCs stained with indicated antibodies. Scale bar 50µM. **D).** Representative IF images of control or pembrolizumab-treated iPDCs stained with CK7 antibody. Scale bar 50µM. **E).** Box plot showing the ratio of CK7+ cells over all cells normalized to control. Analysis was performed 4-6 days post pembrolizumab treatment. Each dot represents an individual image. **F).** Heatmap showing flow cytometry analysis of indicated marker expression in CD8+ T cells, and CD11c+ dendritic cells from iPDCs upon pembrolizumab treatment. Wilcoxon signed rank test ** p< 0.01 *** p<0.001 for (F).

Next, we assessed the presence of source tumor-specific immune cells in OmGel cultures. We performed multi-parameter flow cytometry on freshly-dissociated tumor cells (tumor) and matched PDCs cultured for 4– 7-days and analyzed the data similarly as before (**Fig. 2, A and S2, A**). The results showed that the proportions of the cell types varied between tumor and corresponding PDCs; but this difference was not statistically significant, as tested using paired wilcoxon signed rank test. Importantly, T cells, NK cells and dendritic cells required for eliciting response to immunotherapies were maintained in the cultures (**Fig. 2, B**). Moreover, immunofluorescence staining of selected immune cell markers showed not only the presence of CD45+ immune cells, CD8+ T cells, CD68+ and CD163+ macrophages in the cultures but also spatial interaction between immune cells and CK7+ tumor cells (**Fig. 2, C**). Together, these results validate that HGSC iPDCs preserve source tumor-specific immune cells.

To evaluate the utility of iPDCs for testing immunotherapies, we treated iPDCs with anti-PD1 antibody, pembrolizumab and 3-4 days post-treatment performed single cell image-based analysis to measure tumor cell-specific response, and flow cytometry to evaluate functional states of CD8+ T cells. Our results showed that similar to olaparib response, HRD tumor-derived iPDCs showed reduction in CK7+ tumor cells upon pembrolizumab treatment, which was not observed in case of HRP tumor-derived iPDCs (**Fig. 2, D and E**). Furthermore, flow cytometry analysis revealed, an increase in CD8+ T cell activation markers: Granzyme B (GrzB), IFNγ, and Ki67 upon pembrolizumab treatment, specifically in HRD-iPDCs (**Fig. 2, F**). In summary, these results show that iPDCs enable the investigation of tumor cell-specific growth inhibitory response and immune cell-specific functional responses to immunotherapies.

### iPDCs recapitulate clinical response to olaparib or chemotherapy and reveal drug sensitivities correlating with distinct epithelial cell pathway activities in the resistant tumors

To further establish how iPDC-based drug responses correspond to real-world clinical response, we processed tumors from three patients whose disease had progressed during to olaparib or chemotherapy treatment, and subjected the tumors for genomic, transcriptomic analyses and drug testing on iPDCs (**Fig. 3, A**). In all three patients, sampling was performed in the secondary surgery in case of oligometastatic recurrent disease. While two patients (#C865, #C218) had received chemotherapy and PARPi olaparib before the surgery, the third patient (#C429) was treated with chemotherapy. Interestingly, patient #C865 has continued olaparib maintenance therapy after the surgery, and she is still showing a response with a stable disease (42 months). Conversely, the other two patients, #C218 and #C429, developed progressive disease after the surgery leading to death.

**Fig. 3.**
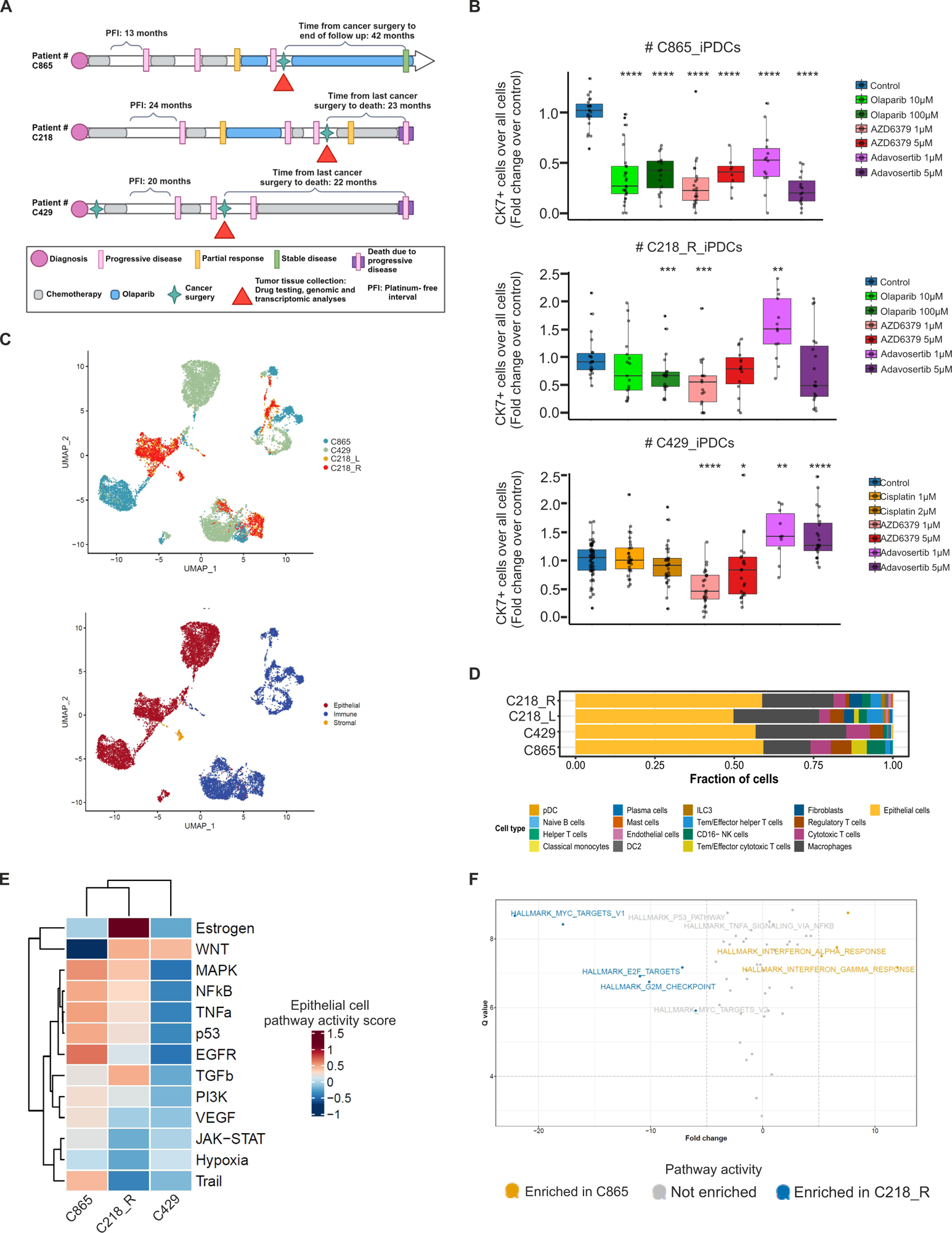
iPDCs recapitulate clinical response to olaparib or chemotherapy and reveal new drug sensitivities. **A).** Schematics representation of clinical course of three patients who were resistant to PARPi or chemotherapy at the time of tumor tissue collection for scRNA seq, and drug testing using iPDCs. **B).** Box plots showing the abundance of ratio of CK7+ cells over all cells, normalized to control upon 3-4 days of indicated drug treatments on the iPDCs of the patients shown in (A). Each dot represents an individual image. **C).** UMAP plot of all cells passing the quality control from scRNAseq data, colored by patient code (top), or by the cell types (bottom) **D).** Box plot showing fractions of cell types across samples shown in (C). **E).** Heatmap of PROGENy scores for epithelial cells from the samples shown in (C). **F).** Scatter plot showing the comparison of pathways enriched in epithelial cells from C865 vs C218_R.

The tumors collected from three patients at indicated time (**Fig. 3, A**) were processed for WES, scRNA-seq, and iPDC establishment for drug testing. The iPDCs from the patients exposed to chemotherapy and olaparib (#C865, #C218_R (right ovary)) were treated with AZD6379 (ATRi), adavosertib (WEE1i), or olaparib (PARPi), while the iPDCs from chemotherapy exposed patient (#C429) were treated with AZD6379, adavosertib, or cisplatin for 3-4 days. At the end of the drug treatment, iPDCs were fixed and stained with anti-CK7 antibody to measure tumor cell abundance as a response to the drug treatment. Our results showed that olaparib, AZD6379, and adavosertib treatment resulted in a significant reduction in CK7+ tumor cells in the iPDCs from olaparib-responder patient (#C865) compared to the control (**Fig. 3, B**). On the other hand, iPDCs from the two resistant patients showed significant response to ATRi, but not to WEE1i. Importantly, consistent with clinical responses, no significant reduction in CK7+ cells were detected following treatment with 10 uM olaparib or cisplatin in iPDCs from olaparib (#C218_R), or chemotherapy (#C429) resistant patients, respectively. Together, these results show that iPDCs recapitulate patient-specific clinical responses to olaparib or chemotherapy.

To gain deeper insights into the molecular features driving drug sensitivity and resistance we performed scRNA-seq and WES on tumors from resistant patients. Uniform Manifold Approximation and Projection (UMAP) plots of all the cells passing the quality control of scRNA-seq data showed patient-specific as well as cell type-specific clusters (**Fig. 3, C and Fig. S3, A**). In addition, epithelial cell and macrophages were the predominant cell types accounting for >70% of all cells identified with scRNA-seq (**Fig. 3, D**). Epithelial cells progeny pathway scores per sample revealed that olaparib resistant tumor (#C218_R) showed enrichment of Estrogen, WNT, and TGFb pathways implicated in resistance to PARPi, in ovarian cancer (*28–30*), while chemotherapy resistant tumor (#C429) was enriched only in WNT signaling. On the contrary, TRAIL (tumor-necrosis factor related apoptosis-inducing ligand) pathway was enriched in the tumor from the olaparib sensitive patient (**Fig. 3, E**). Furthermore, cytotoxic T cells and macrophages from resistant tumors showed distinct activities related to WNT, and JAK-STAT signaling, respectively (**Fig. S3, B and C**). Previous reports show that a sub-set of PARPi resistant clones exhibiting enrichment of G2M Checkpoint, E2F targets are highly resistant to WEE1i (*31*). In line with these results, we found that olaparib and adavosertib resistant sample, #C218_R show enrichment of hallmark of cell proliferation-related pathways such as, G2M Checkpoint, E2F Targets, MYC Targets V1 compared to olaparib sensitive tumor. Multiple mechanisms contributing to PARPi resistance have been identified, including restoration of the HR status, or BRCA1/2 protein expression (*31*). The olaparib resistant patient (#C218) was diagnosed with a germline *BRCA1:c.737delT* mutation. However, somatic alterations analysis of the #C218 resistant tumor through WES revealed a deletion spanning from intron 6 to 7 to the middle of exon 10 on chromosome 17 (chr17:g.43092980-43102933del) both in tumor and iPDCs, causing a potential in frame transcript and restoration of BRCA1 activity. In addition, functional HRD test (*32*) based on RAD51 foci on the formalin fixed tissue samples of these three tumors showed that the olaparib resistant #C218 sample was HRP while other two samples were found to be HRD, supporting the notion of HR restoration in #C218. Together, the results from scRNA-seq and WES showing enrichment of pathways related to cell cycle, proliferation, and drug resistance, combined with restoration of HR proficiency suggest possible mechanisms of therapy resistance leading to progressive disease in olaparib resistant patient #C218. Collectively, our data show that integration of tumor genomic, transcriptomic and drug response analyses from iPDCs enable investigation of drug sensitivity and resistance mechanisms in therapy resistant HGSC patients.

### iPDCs provide a platform for high-throughput drug testing and evaluation of tumor and immune cell-specific responses to immunotherapy combinations

We established a pipeline for testing the feasibility of using iPDCs for high-throughput combinatorial drug treatment as well as for comparative visualization of tumor or immune cell-specific drug response from multiple patients (**Fig. 4, A**). We tested DNA damaging agents such as PARPi, ATRi, WEE1i, along with established immunotherapy agent, anti-PD1 antibody, as well as novel immunotherapies targeting the LPA, or PERK pathway, on 8 individual HGSC-iPDCs cultured in 384 well format (**Table. S, 4**). Each 384 well plate consisted of 54 different conditions comprising 4 concentrations of single agents and 2 concentrations of combination treatments in a replicate of 4 wells. Drugs were dispensed in a randomized manner using a Tecan D300e drug dispenser. As a read out of the treatment response, we employed four channels IF staining followed by high-content imaging to measure single-cell cytotoxic responses from tumor (live / dead CK7+) and immune cells (live / dead CD45+).

**Fig. 4.**
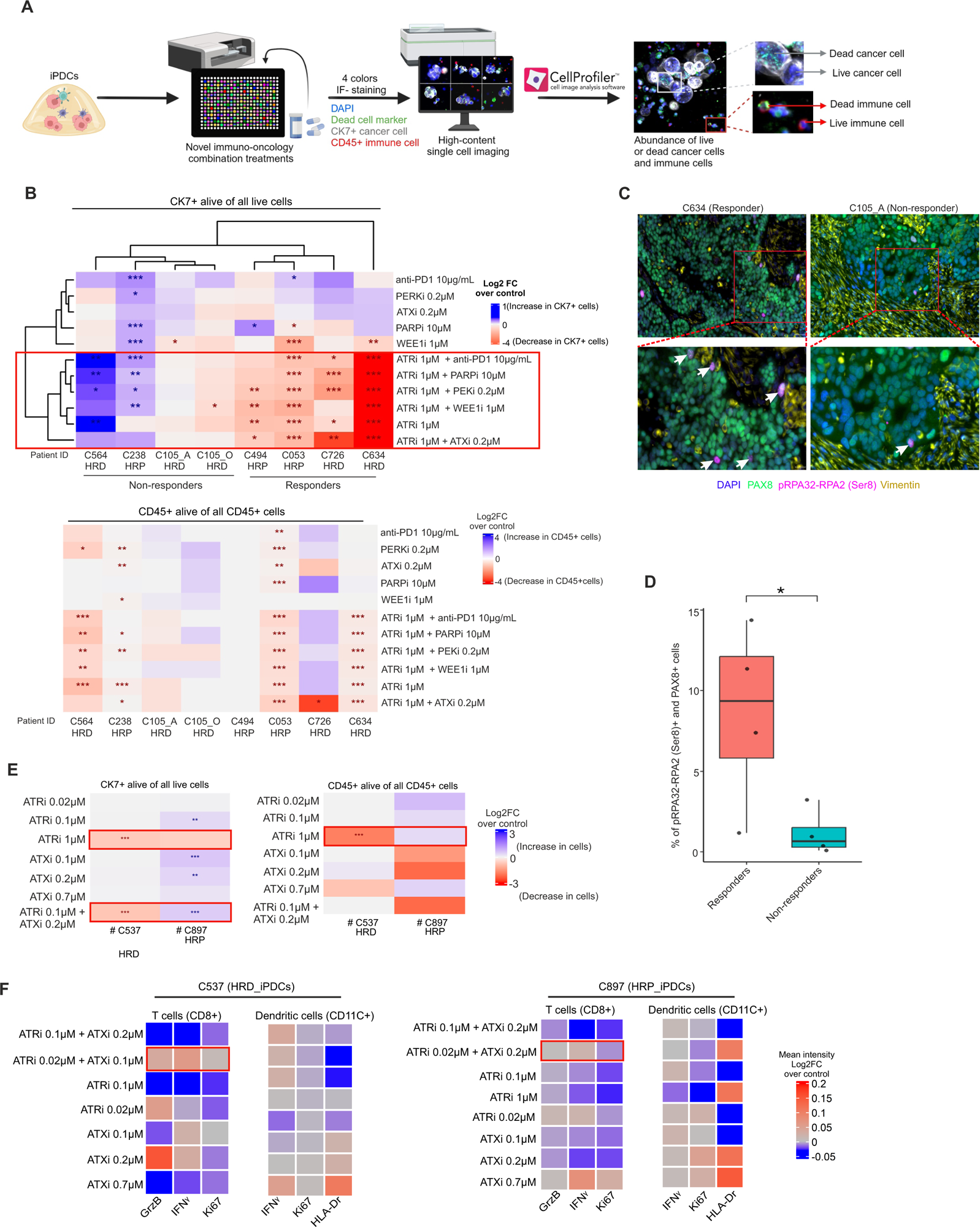
iPDCs provide a plat form for high throughput testing of immuno-oncology agents and evaluation of tumor and immune cell-specific responses to drug treatments. **A).** Schematics showing the workflow for testing immune-oncology agents in 348 well format followed by analysis of tumor and immune cell-specific responses using high-content single cell imaging. **B).** An unsupervised hierarchical clustering showing the Log2FC in live CK7+ tumor cells over all live cells in each treatment condition normalized to control (top), heatmap showing the Log2FC in the live CK45+ immune over all CD45+ cells in each treatment condition normalized to control (bottom). Drug treatments were performed for 4-6 days depending on the sample. **C).** Representative t-CyCIF images on FFPE sections of the tumors from responder (C634) and non-responder (C105_A) to the ATRi combinations, with indicated antibodies. White arrow points towards pRPA32-RPA2 (Ser8)+ and PAX8+ cells. Scale bar, 100µM for the top and 50µM for the bottom panel. **D).** Box plot showing the quantification of the pRPA32-RPA2 (Ser8) staining in PAX8+ cancer cells form the t-CycIF staining on FFPE tumor tissue sections of the samples shown in B). **E).** Heatmaps depicting the Log2FC in the live CK7+ cells over all live cells (top), or live CD45+ cells over all CD45+ cells (bottom) in the indicated samples following different drug treatments for 3 days (C537), or 5 days (C897). Blue asterisks indicate significant increase in live CK7 / CD45+ cells, red asterisks indicate significant decrease in live CK7 / CD45+ cells compared to control. **F).** Heatmaps showing the flow cytometry analysis represented as Log2FC in the mean intensities of the indicated markers in CD8+ T cells or CD11c+ dendritic cells normalized to control from the samples shown in (C) following 3-5 days of treatment with indicated agents. Wilcoxon signed rank test p* < 0.05, p** < 0.01, p*** <0.001 for image analysis. One-sided Mann-Whitney U test p* < 0.05 for quantified data from t-CycIF images.

Unsupervised hierarchical clustering of live CK7+ tumor cells showed that samples clustered based on the drug treatment responses **(Fig. S4, A).** Particularly, in 5/8 samples single agent ATRi or its combination with other agents was effective in reducing the proportion of live CK7+ cells compared to control. Deeper investigation of the significance of this effect revealed that indeed ATRi single and combinatorial treatment with other DNA damaging or immunotherapy agents cause significant tumor cell death in a sample-specific manner (4/8 iPDCs) (**Fig. 4, B**). Notably, this effect was independent of tumor HR status. The analysis on CD45+ immune cell viability revealed that some of the samples showing tumor cell sensitivity to ATRi single and combination treatments also showed decrease in live CD45+ immune cells indicating that the treatment induces toxicity to normal immune cells. However, in two of the responding iPDCs (C053, C634), the anti-tumor effects of ATRi combinations were higher than the cytotoxic effect on immune cells as the fold change in the reduction of live CK7+ cells was higher compared to live CD45+ cells. Furthermore, some of the samples showing sensitivity to ATRi combinations (C494, and C726) showed no significant change in the viability of immune cells, suggesting that the anti-tumor effect is superior to anti-immune cell effect in these samples. Conversely, in two samples (C564, C238) ATRi combinations increased the proportions of live CK7+ cells while decreasing the abundance of live CD45+ immune cells. In addition to ATRi single agent, WEE1i alone showed decrease in tumor cell viability in 3 / 8 samples without significant effects on the immune cell viability. Again, this effect was independent of the tumor HR status **(Fig. 4, B)**. These results suggests that ATRi in combination with other DNA damaging agents: PARPi, or WEE1i, as well as immunotherapies: anti-PD1, ATXi, or PERKi is effective against a sub-set of HGSC iPDCs. Importantly, we show that assessment of immune cell-specific death response in addition to tumor cell-specific response, is necessary to evaluate the benefit of the immuno-oncology combinations.

Increase in replication stress markers such as phosphorylated replication stress protein A (RPA) is known to be predictive biomarker for sensitivity to ATRi (*33*). To evaluate if this is also reflected in the patient’s tumors, we performed tissue cyclic multiplex immunofluorescence (t-CycIF) (*34*) using antibodies against PAX8 (tumor cells), vimentin and replication stress marker pRPA32-RPA2(Ser8) on the corresponding formalin-fixed paraffin embedded (FFPE) tumor tissues from the same patients shown in (**Fig. 4, B**). Single cell analysis of t-CycIF images for PAX8 positive cancer cells expressing pRPA32-RPA2(Ser8) revealed that the tumors responding to ATRi combinations had significantly higher fraction of pRPA32-RPA2(Ser8) + and PAX8+ cells compared to non-responders (**Fig. 4, C-D**). These results indicate that our platform allows screening of novel combinatorial treatments on iPDCs along with investigation of tumor tissue-based predictive biomarkers.

The results from our high-throughput combination treatments provide insights into the tumor and immune cell-specific effects of single agent ATRi and in combinations with other agents. Since our results suggested cytotoxic effects of ATRi combinations on immune cells in a sample-specific manner, we tested if lowering the concentrations of ATRi had beneficial effects on immune cells yet cytotoxic for the cancer cells. We selected ATXi for further validation and functional analysis in combination with ATRi as this agent is known to induce CD8+ T cell activation via inducing interferon signaling in dendritic cells (*9*). To further analyze the functional effects of ATRi and its combinations on the different immune cell subtypes, we evaluated tumor and immune cell-specific death responses, as well as CD8+ T cell and CD11c+ dendritic cell functional states using flow cytometry. Consistent with previous results, we found a significant decrease in tumor and immune cell viability following treatment with 1µM ATRi in one of the two tested tumors. However, neither the lower concentrations of ATRi or single agent ATXi caused significant reduction in tumor or immune cell viability. Interestingly, combination of 0.1µM ATRi with 0.2µM ATXi had an opposite effect on the two tumors: decreasing the proportion of live CK7+ tumor cells in C537_iPDCs (responder) while increasing the proportion of live CK7+ tumor cells in C897_iPDCs (non-responder). In addition, the responder sample did not show significant reduction in immune cell viability upon treatment with 0.1µM ATRi + 0.2µM ATXi, while the non-responder showed a trend towards a decrease in live CD45+ immune cells (**Fig. 4, E**). Furthermore, flow cytometry analysis showed that upon 0.02µM ATRi and 0.2µM ATXi combination treatment, CD8+ T cell activation markers GrzB, and interferon gamma (IFNγ) were increased in CD8+ T cells, and IFNγ was increased in CD11c+ dendritic cells from responder but not in the non-responder (**Fig. 4, F**). Together, these results show that HGSC-iPDCs support cytotoxic and functional response analysis from tumor cells and immune cells, respectively upon treatment with novel immuno-oncology agents.

## DISCUSSION

A variety of pre-clinical models of HGSC have been established for disease modelling and drug testing. Some of these include two-dimensional (2D) cell cultures, three-dimensional (3D) spheroids and organoids, tissue slices, microfluidics (organ-on chip), and mouse models (*35*). These models represent different levels of biological complexities found in patient tumors: 2D and 3D models depict sub-cellular and cellular complexities, tissue slices, and microfluidics mimic tissue and organ level complexities, mouse models elucidate organism-level understanding of the disease progression. While 2D and 3D models allow long-term cultivation and selective growth of cancer cells, they lack the TME and are vulnerable to genetic changes over the time. On the other hand, tumor tissue slices, and microfluidics are short-term cultures are not suitable for high-throughput drug response evaluation. Therefore, there is an unmet need for a pre-clinical model of HGSC that retains both genetic complexity and the microenvironment of the patient tumors yet is amenable for high throughput testing of immuno-oncology agents.

Here we established a novel patient-derived immunocompetent culture of HGSC tumors using patient derived OmGel matrix, while preserving the native tumor-infiltrating immune cells. Previous studies on HGSC organoid cultures have utilized mouse sarcoma derived BME matrix with an average of 89 days to establish the cultures, with the majority of the successful cultures from ascites (*36*). Our result demonstrating that the same tumor derived cells cultured in OmGel or BME show significantly improved growth in OmGel within 7 days of the culture suggest that OmGel provides a growth environment analogous to the HGSC metastatic tumors. As opposed to long-term organoid cultures of HGSC, iPDCs are short-term cultures lasting up to one week without the ability to passage over time. This is an inherent limitation of any relevant immuno-oncology *ex vivo* models that can sustain the viability and functional states of the diverse immune cell repertoire.

The proteomic analysis of OmGel by Neilson et al., report that there are no detectable levels of cytokines and growth factors in OmGel, suggesting that OmGel on its own does not alter the immune compartment of the iPDCs (*21*). In line with this we show that neither dispase digestion nor OmGel matrix alter the immune cells indicating that all the immune cells from the source tumors are faithfully recapitulated in the iPDCs, thereby making them an ideal model for the investigation of immunotherapy agents. Although the evidence from clinical trials indicates unfavorable response to ICIs in HGSC, these studies were conducted on patients pre-treated typically with multiple lines of chemotherapy, and platinum sensitivity positively correlated with response to single agent ICIs, and HRD biomarker associated with the efficacy to combined PARP plus PD-1 inhibition 5. However, studies on of ICIs on treatment naive patients, especially stratified by HRD status have not been reported. Here we provide novel insights that iPDCs established from treatment naive tumors show HRD-specific response to pembrolizumab both in terms of reduction in tumor cells and activation of CD8+ T cells. Prior to clinical investigation, validation of this finding on a bigger cohort of iPDCs is warranted.

Apart from the ability of iPDCs to evaluate response to immunotherapy agents, our results highlight that iPDCs mimic the clinical responses to PARPi, and chemotherapy treatments thus validating the utility of iPDCs for testing chemotherapy and targeted therapies. In line with the clinical responses to PARPi, we found 2/3 HRD-iPDCs showed responses to PARPi Olaparib. Combining the drug response from treatment resistant iPDCs with pathway analysis from scRNA-seq and mutation analysis from WES suggested possible mechanism of resistance to PARPi. These include enrichment of cell cycle, G2M checkpoint, WNT, and TGFb pathways in PARPi resistant tumor, an acquired *BRCA1* mutation likely leading to restoration of HR function. These findings support the concept of integrating drug sensitivity and resistance profiling of iPDCs with molecular features from the corresponding tumors to reveal mechanism behind treatment responses to enable implementation of more effective treatment strategies.

Thus far, high-throughput drug response profiling in HGSC has been performed predominantly on cancer cell cultures or organoids using compounds targeting oncogenic drivers. Herein we have established a patient-derived pre-clinical model and analysis pipelines that facilitate the investigation of immuno-oncology combinations in a high-throughput manner enabling the analysis of treatment responses to over 50 different conditions on tumors from multiple patients. Moreover, while previous studies have mainly involved measurement of metabolic markers independent of the cell types, our single-cell image-based readout uniquely captures tumor and immune cell specific functional states. Although combination of ATRi with other DNA damaging agents such as PARPi, or WEE1i have been clinically tested in ovarian cancer (*37, 38*), the combination of ATRi with immunotherapy has not been investigated before. For the first time, we report that combination of ATRi with both investigational (ATXi, and PERKi), and established (anti-PD1 antibody) immunotherapies show patient-specific effects on tumor and immune cell viability. Furthermore, we found that higher concentrations (1µM or above) of ATRi are likely toxic as they caused both cancer and immune cell death in a subset of iPDCs. However, our results using lower concentrations of ATRi (0.1µM) in combination with ATXi showed cancer cell-specific death without affecting immune cell viability in a sample-dependent manner. Importantly, tumor cell-death responses to this combination aligned with CD8+ T cell and dendritic cell activation status. Together these results suggest that iPDCs are suitable for screening immuno-oncology compounds to identify promising drug candidates as well as for deeper analysis of functional response from the immune cells. Furthermore, the results from t-CycIF corroborate previous studies showing replication stress as a biomarker of sensitivity to ATRi and suggest that this approach aid in the investigation tissue-based biomarkers for potential combination therapies identified on iPDCs. Our data from iPDCs highlight that integration of high-content single cell image analysis with immune cell functional states from flow cytometry enable assessment of tumor cell-specific cytotoxic and immune cell-specific functional responses for stratification into responders and non-responders. Together, our novel HGSC patient-derived immunocompetent cultures in physiologically relevant matrix combined with patient-specific tumor infiltrating immune cells facilitates high-throughput testing of novel immuno-oncology agents and personalized combination therapies in HGSC.

## MATERIALS AND METHODS

### Patient material and clinical data

All experiments involving patient materials were performed in accordance with the ethical standards from the 1975 Declaration of Helsinki. Tissue samples were collected as a part of ONCOSYS-OVA study (clinical trial number: NCT06117384) from HGSC patients undergoing primary debulking surgery (PDS) (n=36) or secondary surgery (n=3) at Helsinki University Hospital (HUH). Each patient gave written informed consent and the use of fresh tissue, blood, and formalin-fixed paraffin embedded (FFPE) HGSC clinical material was approved by the Ethics Committee of Women’s Clinic, HUH and Helsinki Biobank.

### Tumor tissue processing

In this study, we processed HGSC tumors from 39 patients, depending on the availability tumor tissues were subjected to different experimental procedures listed in **Table. S, 1**. Freshly resected tumor tissue was cut into multiple pieces (approximately 100 mm^3^ per piece). Three to six pieces were snap frozen in liquid nitrogen for DNA, and RNA, isolation, one piece was fixed in 10% formalin for histopathology, and t-CycIF analysis. Depending on the availability100 mm^3^-2000 mm^3^ cm tumor tissue piece was minced into smaller pieces and digested with 1x Collagenase & Hyaluronidase (# 7912, Stem Cell Technologies), or 1U/mL Dispase II (# D4693, Sigma/Merck), in ADF+++ medium (DMEM F12, (# 11320-074 Thermo Scientific) supplemented with GlutaMAX™-I (100X) (#35050-061, GibcoLifeTechnologies), Penicillin-Streptomycin (10,000 U/ml), (#15140122 GibcoLifeTechnologies), and HEPES, (# H0887, Sigma)), and 50 U/ mL DNAse (# M6101, Promega) at 37°C for 1 hr rotating incubator. Digested tissue was filtered through 70µM strainer and centrifuged at 1000 RPM for 10 min at +4°C. The pellet was subjected to RBC lysis for 5 min at room temperature (RT) followed by dilution in AdF+++ medium and centrifuged at 1000 RPM for 10 min at +4°C. Cells in the resulting final pellet was counted with trypan blue using LUNA-FL™ Dual Fluorescence Cell Counter.

### Viability and Immunofluorescence analysis on the dissociated cells

Enzymatically dissociated cells were plated on to Shandon™ Multi-Spot Slides (#9991090, ThermoFisher Scientific) at a concentration of 30000 cells / 3µL and allowed to air dry at RT. Cells were stained with LIVE/DEAD™ Viability/Cytotoxicity Kit, for mammalian cells (# L3224, Thermo Fisher Scientific), and incubated at 37°C for 30 min, or cells were fixed with 4% paraformaldehyde for 10 min at RT. Followed by blocking with 2% TritonX-100+3%BSA in 1xPBS for 30 min at room temperature, and staining with Anti-Cytokeratin 7 antibody [EPR17078] (Alexa Fluor® 555, ab209601), and ati-CD45 [HI30] (Alexa Fluor® 647, #304018 Bio Legend) for 1hr at RT. Images were acquired using Nikon Slide express 2 microscope, followed by image analysis using Cell Profiler. The quantified data was plotted using GraphPad prism.

### Quantitative real-time PCR *(RT-qPCR)*

RNA was extracted from the cells dissociated with C&H or dispase or from fresh frozen tissues using RNeasy plus mini kit (#74134, Qiagen). 500 ng of RNA from each sample was used for cDNA synthesis using High-capacity RNA-to-cDNA kit (#4387406, Thermo Fisher Scientific). qPCR mixture was prepared using PowerUp™ SYBR™ Green Master Mix for qPCR (#A25741, Thermo Fisher Scientific), and ran using BioRad machine CFX 384. The quantified data was plotted using GraphPad prism.

### Establishment of patient-derived cultures

Patient-derived OmGel was prepared as previously described (*21*). HGSC tumor-dissociated cells were mixed with Cultrex RGF Basement Membrane Extract (BME), Type 2 (# 3533-010-02 R&D systems), or 2.5 mg/mL OmGel at a concentration of 1000 cells / µL and plated as a droplet in 48-well or 96 well plates. OmGel mixture was prepared as follows: appropriate volume of cell suspension was mixed with, 2.5 mg/mL OmGel, 33.3 ug/mL aprotinin (# A6279 Sigma), 0.3 U thrombin (#T6884 Sigma), and 0.5 µg/mL of fibrinogen (# 341578 Sigma). Addition of fibrinogen initiates solidification of the gel, hence it was added just before plating the suspension. Droplets containing BME or OmGel resuspended cells were allowed to solidify at 37 ^0^C incubator for 1 hr. iPDC culture medium II (Supplementary Methods. Table 1) supplemented with growth factors and cytokines was added on top of the droplets and cultured for 4-10 days depending on the sample.

**TABLE 1:** Composition of iPDC growth medium. Once prepared, medium I was used for up to one month. The medium II supplemented with cytokines is freshly prepared prior to setting up of iPDC cultures.

| Name | Name | Stock Concentration | Working Concentration | Company/ Catalogue |
| --- | --- | --- | --- | --- |
| Medium I | Advanced DMEM/F12 |  |  | Thermo Scientific/ #12634-010 |
|  | Primocin | 50 mg/ml | 100 µg/ml | Invivogen / #ANTPM1 |
|  | NEAA | 100x | 1x | Gibco/ #11140035 |
|  | Sodium Pyruvate | 100x | 1x | Gibco/ #11360-039 |
|  | L-Glutamine | 100x | 1x | Gibco Life Technologies/ #35050-061 |
|  | HEPES | 100x | 1x | Sigma Merk/ #H0887-100ML |
|  | N-Acetylcysteine | 100 mM | 1 mM | Sigma/ #A9165-100G |
|  | Nicotinamide | 100 mM | 5 mM | Sigma/ #N0636-500G |
|  | Supplement B-27 | 50 x | 1X | ThermoFischer/ #17504044 |
|  | FGF-10 | 100 µg/mL | 10 ng/mL | Peprtech/ #100-26 |
|  | basic FGF (FGF4) | 100 µg/mL | 10 ng/mL | Peprtech/ #AF-100-18B |
|  | A-83-01 | 10 mM | 0.5 µM | Sigma/ #SML0788-5MG |
|  | β-estradiol | 10 mM | 10 nM | Sigma/ #E2758-1G |
|  | Neuregulin-1 / Heregulin | 6.6 uM / 50 ug/ mL | 5 nM | Peprtech/ #100-03 |
|  | EGF | 100 µg/mL | 5 ng/mL | Peprtech/ #AF-100-15 |
|  | Hydrocortisone | 1 mg/mL | 500 ng/mL | Sigma/ #H0888 |
|  | Y-27632 / ROCK inhibitor | 5 mM | 5 µM | AbMole Biosciences/ #M1817 |
| Medium II<br>(Medium I + cytokines) | IL2 | 66 X E6 IU/MI | 500 IU/mL | Peprtech/ #200-02 |
|  | IL15 | 10 µg/ml | 10 ng/ml | Peprtech/ #200-15 |
|  | IL7 | 10 µg/ml | 10 ng/ml | Peprtech/ #200-07 |

### Quantification of number of iPDCs per sample

On day 4-10 of the BME or OmGel cultures, bright-field images from 4 wells were taken at 10x magnification, and the number of cell clusters >3000 µm2 was counted using ImageJ-Fiji. The data was plotted using GraphPad prism.

### Whole-exome sequencing

iPDCs were harvested using Cultrex Organoid Harvesting solution; # 3700-100-01), and DNA was extracted using AllPrep DNA/RNA Mini Kit (Qiagen). 50ng of gDNA from blood (germline, n=11), fresh frozen tumor (n=13), FFPE tumor tissue (n=2), and iPDC (n=10) samples were subjected to WES. WES libraries were prepared using Twist Comprehensive Exome according to the manufacturer’s instructions. This process involved the enrichment of exonic regions, capturing a comprehensive representation of the protein-coding regions of the human genome. Prepared libraries were subjected to high throughput sequencing using Illumina NovaSeq 6000 system using S4 flow cell (Illumina, San Diego, CA, USA) and v1.5 chemistry. DNA samples were subjected to quality control, alignment to the reference human genome, deduplication, estimation of cross-sample contamination, and discovery of genetic variants. We assessed organoid-tissue mutations concordance and Variant allele frequency (VAF) correlation using 10 organoid-tissue pairs. Details of the data processing and analyses can be found in the Supplementary Methods.

### Flow cytometry and data analysis

Tumor tissue or iPDCs were dissociated using Dispase II (# D4693, Sigma/Merck), and the cells were stained with two different antibody panels, immune cell panel for comparison of dissociation methods consisted of following antibodies: anti-CD14 PE (18D11, #21620144, ImmunoTools), anti-HLA-Dr PerCP-CY5.5 (G46-6, #552764 BD), anti-CD8 PE-Cy7 (RPA-T8, #557746, BD), anti-CD4 APC (MEM,241 #21270046, ImmunoTools), anti-CD56 AF700 (B159, #557919, BD), anti-Epcam APC/Fire 750 (9C4, #324233, Nordic BioSite), anti-CD45 BV421 (HI30, #563880, BD), anti-CD16 BV510 (3G8, #563829 BD), anti-CD19 BV605 (HIB19, #302243, BD), anti-CD11c, BV11c (B-ly6, #563130, FisherScientific), anti-CD3 BV786 (CD3 SK7, #563800, BD), dead cell marker (DCM) FITC (#L2301,Thermo Fisher Scientific). Analysis of the functional states of the immune cells consisted of: anti-CD56 FITC (MEM-188, #21270563, ImmunoTools), anti-CD4 PE-CF594 (RPA-T4, #562281, BD), anti-Ki67 PE-CY7 (B56, #561283, BD), anti-Granzyme APC (GB11, #GRB05, Thermo Fisher Scientific), anti-IFN γ AF700 (B27, #557995, BD), DCM BV510 (#L34965 Thermo Fisher Scientific), and the remaining antibodies were same as in the immune cell profiling panel. Samples were run using BD LSRFortessa™ Cell Analyzer. From the flow cytometry data, DCM^-^ EpCAM^-^ cells were gated using FlowJo, and exported as csv files for further analysis using CYTO (Figure S2A) (*27*). The proportions of the cell types from the clusters generated in CYTO were plotted using R-script.

### Drug treatment and immunofluorescence staining

Dissociated patient-derived cells were embedded as 1000 cells / µL in 25 µL of OmGel matrix on a CellCarrier 384 Ultra microplates, PerkinElmer, or in 10 µL of OmGel matrix on a Cell Culture 96 well F-bottom microplate, Greiner. Drops were left to solidify for 1 hr at +37°C before adding iPDC-growth medium. Next day, cultures were treated with olaparib (PARPi, AZD2281; SelleckChem), pembrolizumab (anti-PD-1, Selleckchem), ziritaxestat (ATXi, GLPG1690; MedChemExpress), AMG PERK44 (PERKi; MedChemExpress), Adavosertib (WEE1i, MK-1775; SelleckChem) or berzosertib (ATRi, VE-822; SelleckChem) as single agents or as combinations using TECAN D300e Digital Dispenser (TECAN) in a randomized manner. Dimethyl sulfoxide (DMSO) was used as a control. Chemo- and PARPi-resistant patient-derived iPDCs were cultured on a Shandon Multi-Spot slides (#9991090, ThermoScientific) and treated with cisplatin (S1166; SelleckChem) or Olaparib, WEE1i, or ATRi. Drug treatment was carried out for 3-6 days, depending on the growth of iPDCs, as evaluated by phase-contrast microscopy. At the end of drug treatment, cells were stained with live-or-dye stain following manufactureŕs protocol (#32004, Biotium) before fixing with 4% paraformaldehyde for 1hr at RT. Wells were washed 3x with 1x PBS followed by permeabilization and blocking with freshly prepared 3% bovine serum albumin and 0.3% Triton-100 in 1x PBS for 1hr at RT. After removing the blocking buffer, cells were incubated with antibody cocktail containing Anti-Cytokeratin 7 antibody [EPR17078] (Alexa Fluor® 555 conjugated, ab209601), and ati-CD45 [HI30] (Alexa Fluor® 647 conjugated, #304018 Bio Legend), and Hoechst (#3342, Thermo Fisher Scientific) in the blocking buffer overnight at +4^0^C. Next day wells were carefully washed 3 times with 1x PBS to remove all the TritonX-100 and unbound antibody mix. Wells were left with 50 µL of PBS for imaging.

### High-content single-cell imaging and data analysis

Images of the drug-treated samples (10-17 images per well) were acquired using Opera Phenix High Content Screening System (PerkinElmer) with 40x objective and Andor Zyla sCMOS camera with Harmony High Content Imaging and Analysis Software (PerkinElmer). Raw image processing was done with BIAS (Biology Image Analysis Software). Cell segmentation and quantification of tumor (CK7+), immune cells (CD45+), and dead cells (DCM+), was performed using CellProfiler. Sample-specific threshold was set for CK7 integrated intensity values from the membrane, CD45 mean intensity values from the membrane, and mean intensity values from the nucleus of DCM per sample for further analyses.

The data from CellProfiler was graphically represented as the abundance of live tumor (DCM-, CK7+) and live immune cells (DCM-, CD45+) per treatment and per patient sample using RStudio. Outlier images which contained fewer than five cells, no CK7+ cells, *and* no CD45+ cells were filtered out. Images which contained an excessive number of cells, greater than three standard deviations above the mean, were also excluded. The data was visualized using a ratio of CK7 alive over all alive cells (tumor cell abundance), CD45 positive alive cells over all cells (immune cell abundance). The median tumor and immune abundance proportions per treatment were normalized by the control and the log2 fold change was plotted in the heatmap.

### Single-cell RNA-sequencing and data analysis

Dispase-dissociated single cell suspensions were washed in 1X PBS + 0.04% BSA solution and cell viability was assessed on an automated cell counter. scRNA-seq libraries were prepared with Chromium NextGEM Single Cell 3’ reagent kit (10x Genomics) and sequenced on a NovaSeq 6000 instrument (Sequencing Unit of Institute of Molecular Medicine Finland, Finland). scRNA-seq data preprocessing was done as described previously (*23*). Briefly, the FASTQ files were processed with the Cell Ranger pipeline v.6.0.1 (10x Genomics) to perform sample demultiplexing, alignment, barcode processing, and UMI quantification. The reference index was built upon the GRCh38.d1.vd1 reference genome with GENCODE v25 annotation. The count matrices of four samples were loaded into Seurat R package (v.4.3.0) and merged, and themiQC algorithm (*39*) was used for quality control. Cells were assigned to the low-resolution cell types (epithelial tumor cells, stromal cells, and immune cells) based on the expression of marker genes as described previously (*23*). To assign high-resolution immune cell types, the celltypist algorithm was used with the model “Immune_All_Low”.

The decoupleR R package (v.2.7.1) (*40*) was used to infer biological activities of 14 signaling pathways available in PROGENy (*41*). For each sample, pathway activities across cell types were inferred by running the *run_ulm* function with default parameters, and aggregated pathway activities were visualized on a heatmap.

### tCycIF staining and image processing

Formalin-fixed paraffin-embedded (FFPE) tumor tissue sections from eight PDS patient samples were sequentially stained with validated antibodies (Supplementary Methods Table 3) and scanned using RareCyte Cyte Finder as outlined in the t-CycIF protocol (*34*). Images from each cycle of staining were stitched and registered to create one high-plex image using the ASHLAR algorithm (*42*). Nuclear segmentation was achieved with the StarDist method by identifying positive nuclear Hoechst staining (*43*). The mean fluorescence intensity for all markers expressed within each cell was quantified using an in-house python script. In total 6,784,375 cells were identified from which 2,381,181 were annotated as cancer cells.

**TABLE 2:**
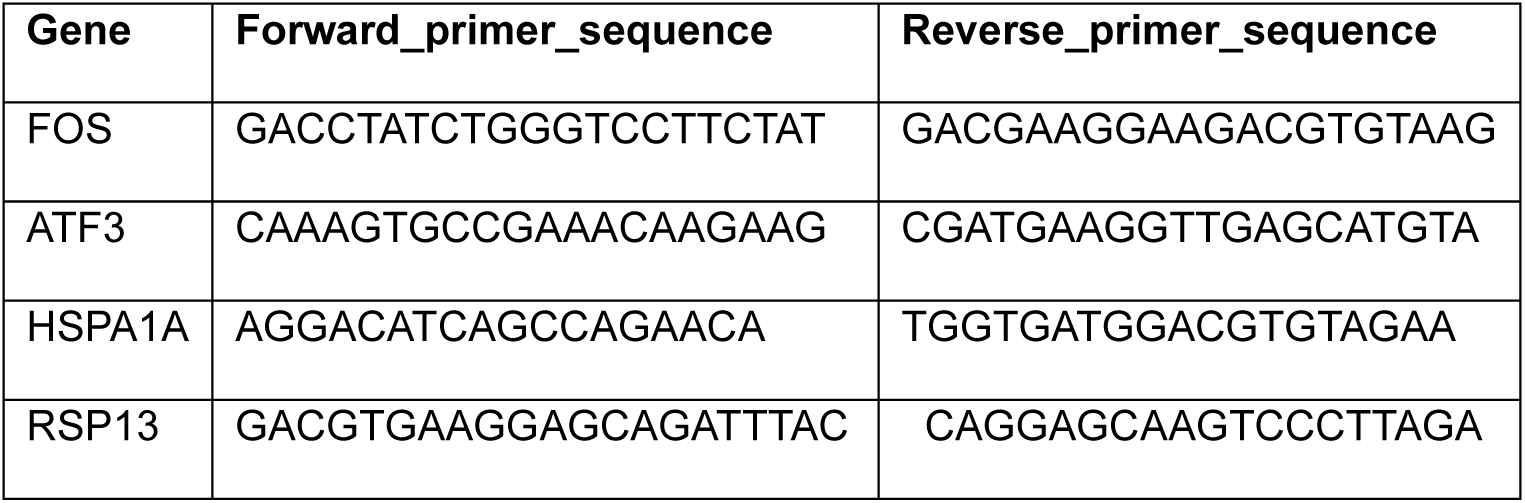
Primers used for the qPCR analysis.

**TABLE 3:** Antibodies used in t-CycIF panel.

| Antibody | Fluorochrome | Company | Cat number | Clone | Concentration | Target |
| --- | --- | --- | --- | --- | --- | --- |
| Hoechst_1 | 361 | Invitrogen | H3570 | *blank* | 10000 | Nucleus |
| Granzyme B | unconj (647) | Dako | M7235 | GrB-7 | 50 | Activated Cytotoxic T and NK cells |
| CD68 | 488 | CST | 24850S | D4B9C | 100 | Macrophages |
| Phospho-Stat1 (Y701) | 555 | CST | 8183S | 58D6 | 100 | Activated INF-γ pathway + cytotoxic T and NK cells |
| CD8a | 660 | Invitrogen | 50-0008-80 | AMC908 | 150 | Cytotoxic T-cells |
| Vimentin | 750 | CST | 9855S | D21H3 | 100 | Stromal cell marker |
| Phospho-RPA32/RPA2(Ser8) | 488 | CST | 31912S | E5A2F | 100 | RPA (replication stress and DNA damage) |
| CD3D | 555 | Abcam | ab208514 | EP4426 | 100 | Immue cell Infiltration |
| PD1 | 647 | Abcam | ab201825 | EPR4877(2) | 150 | Activated T and B cells |
| CD4 | 488 | RnD systems | FAB8165G | polyclonal | 150 | Helper T-cells |
| CD11C | 555 | CST | 77882BC | D3V1E | 150 | Dendritic cells |
| Pax8 | 488 | Abcam | ab214955 | EPR18715 | 150 | HGSC (carcinoma) |
| FoxP3 | 555 | Invitrogen | 41-4777-80 | 236A/E7 | 100 | Regulatory T cells |
| HLA-DPB1 | 647 | Abcam | ab201347 | EPR11226 | 400 | HLA II |
| CK7 | 555 | Abcam | ab209601 | EPR17078 | 200 | Tumor |
| gH2Ax | 647 | BioLegend | 613407 | 2F3 | 150 | DNA Damage |

To compare RPA32 phosphorylation in tumor cells between treatment responders and non-responders, PAX8 and pRPA32/RPA2(Ser8) positive cells were annotated by setting unique intensity thresholds for each sample based on visual inspection in Napari (https://github.com/napari/napari). False positives resulting from artifacts such as folds in the tissue were excluded by setting an upper limit threshold. The proportion of PAX8+ and pRPA32/RPA2(Ser8)+ over PAX8+ cells was calculated per sample and the Mann-Whitney U test was performed to determine statistical significance.

### Statistical analysis

Paired Wilcoxon signed-rank test was applied for comparison of organoid numbers between BME and OmeGel. Pearson correlation was employed for correlation analysis of the WES data. A p-value threshold of 0.05 was set to determine significant statistical comparisons. Paired Wilcoxon signed-rank test was used for comparison of immune cell proportions between C&H vs dispase, BME vs OmGel, and tumor vs iPDCs. Wilcoxon signed-rank test was used to compare the proportions of CK7+ tumor or CD45+ immune cells in each treatment against the control. Mann-Whitney U test was applied for quantified data from t-CyCIF images.

## ACKNOWLEDGEMENTS

This study was co-funded by the European Union (ERC, SPACE 101076096), Sigrid Jusélius Foundation, Cancer Society of Finland, Research council of Finland (grant numbers 1339805, 350396, 351196), iCAN Digital Precision Cancer Medicine. We thank Institute for Molecular Medicine Finland (FIMM) Technology Centre, and Single-Cell Analytics unit, University of Helsinki, for NGS library preparation, WES sequencing, and single cell RNA sequencing. We also thank FIMM High content imaging and analysis unit (FIMM-HCA) for scanning the fluorescent images, and FIMM digital microscopy and molecular pathology unit for tissue processing and scanning the HE images. We also thank Prof. Sara Zanivan for advising with omentum gel-related experiments, and Fernando Perez for valuable advice on the analysis of high-content imaging data.

## AUTHOR CONTRIBUTIONS

A.S.N. designed and performed the experiments, analyzed the data, and wrote the paper. M.S, E.P.A., K.N., S.P performed the experiments. M.S., A.Si., E.P.E., I.N, J.B., G.M., S.P., A.Sa, analyzed the data. A.K., J.T., R.K.K., U.M.H., provided patient samples. A.Vi., performed pathological analysis. S.H., A.Vä., J.T., E.K., L.K. provided the resources and supported data analysis. T.S designed the OmGel preparations and supported the data analysis. All authors contributed to the writing and editing of the manuscript. A.F conceived and supervised the study and wrote the paper.

## DATA AND MATERIALS AVAILABILITY

The iPDC sequencing data, and scRNA-seq data from the resistant tumors will be available in public databases upon publication. Raw images of the iPDC drug treatments will be available in SYNAPSE upon publication. All the codes related to data visualization will be available in GitHub upon publication.

## SUPPLEMENTARY MATERIALS AND METHODS

### WES data processing and analysis

Bioinformatic processing of DNA samples involved several steps, including quality control, alignment to the reference human genome, deduplication, cross-sample contamination estimation, and variant discovery. Sequenced DNA reads underwent quality control and trimming steps using FastQC (*44*) and Trimmomatic tools (*45*). Subsequently, the high-quality reads were aligned to the reference human genome GRCh38.d1.vd1 using BWA-MEM (*46*) with default parameters, subjected to deduplication with Picard tool and base quality recalibration using the Genome Analysis Toolkit (GATK) version 4.1.9.0 (*47*). Cross-sample contamination estimation was conducted using GATK 4.1.9.0, with a contamination estimation threshold established at 15%.

### Mutation calling

Somatic short variants were called through the collective analysis of multiple tumor samples against a singular matched normal for each patient, using GATK 4.1.4.1 Mutect2 according to established best practices. We used a Panel of Normal (PoN) generated with 181 normal samples from the prospective, longitudinal, multiregion observational study DECIDER (Multi-Layer Data to Improve Diagnosis, Predict Therapy Resistance and Suggest Targeted Therapies in HGSOC; ClinicalTrials.gov identifier NCT04846933) and 99 TCGA normals. A detailed description of PoN creation can be found in Lahtinen et al (*48*). Subsequently, GATK FilterMutectCalls was used for variant filtration, keeping only those variants that successfully passing all filters. Variant allele frequencies (VAF) were computed based on the read depths for reference and alternate alleles (AD field). We annotated the using ANNOVAR 20191024 (*49*).

### iPDC-tissue concordance evaluation

To assess the alignment between iPDC and tissue samples, we calculated tissue-based concordance (Ct) by determining the ratio of shared mutations in both iPDC (i) and tissue samples (t) to mutations found in the tissue (t) Ct = (Mi ∩ Mt) / Mt. Concordance analysis was conducted for iPDC-tissue pairs, with each pair consisting of iPDC and tissue samples derived from the same patient and organ. In total, 10 iPDC-tissue pairs were analyzed.

Like iPDC-tissue concordance, we computed organoid-tissue Variant Allele Frequency (VAF) correlation for 10 pairs composed of iPDC and tissue samples from the same patient.

**Table S1.**
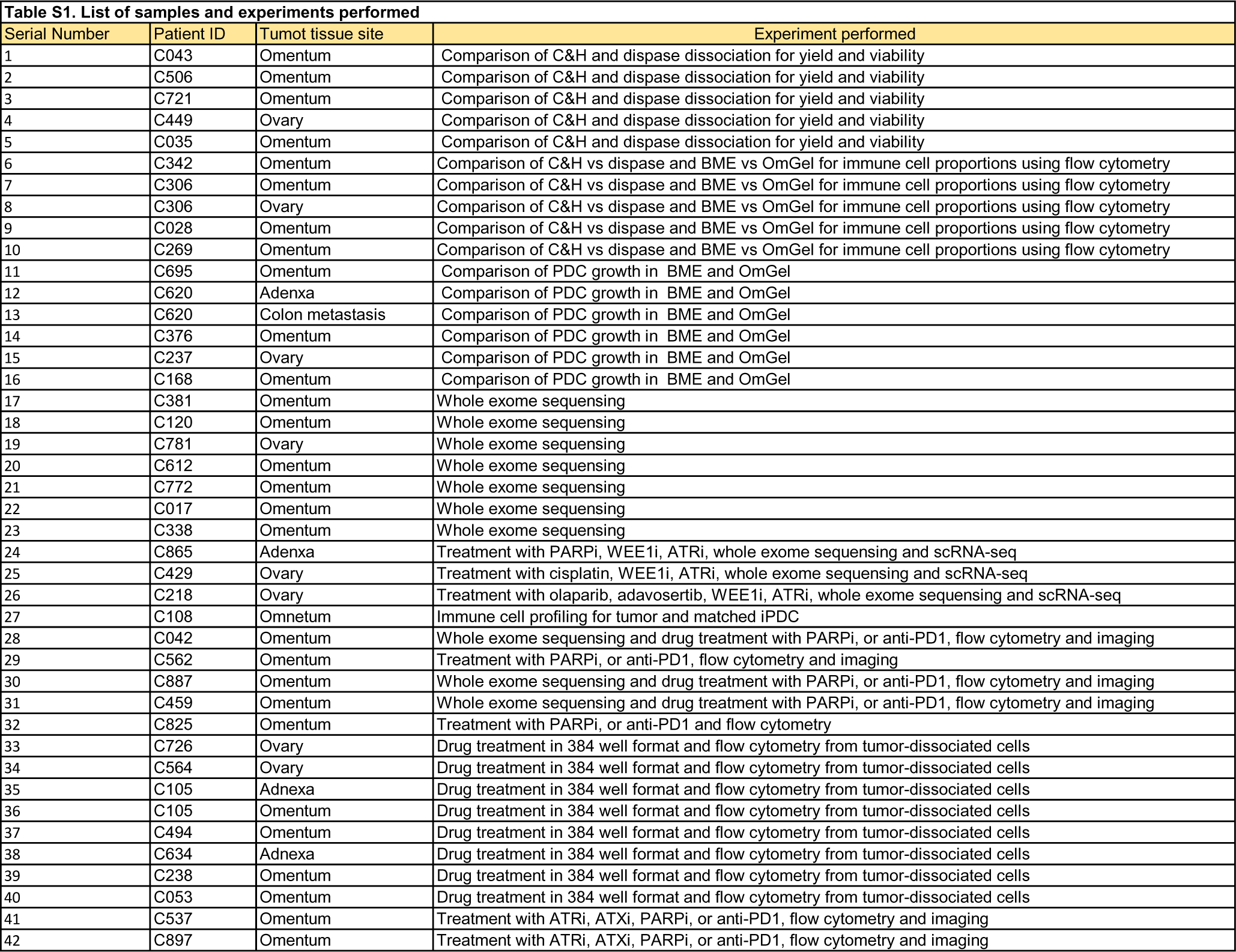
List of samples and experiments performed.

**Table. S2.**
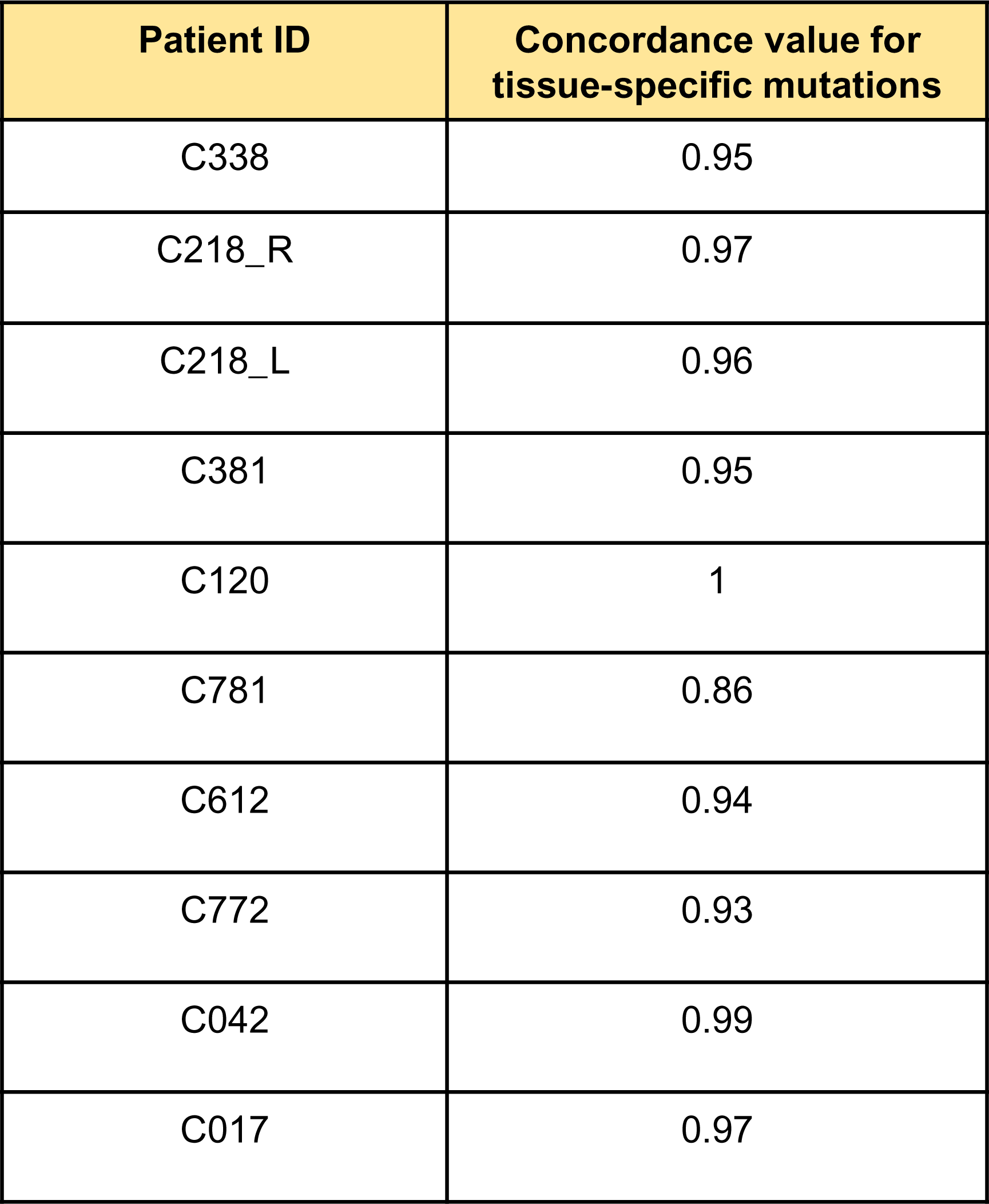
Concordance of tissue specific mutations for individual tumor - iPDC pairs.

| **Patient ID** | **Concordance value for**<br>**tissue-specific mutations** |
| --- | --- |
| C338 | 0.95 |
| C218_R | 0.97 |
| C218_L | 0.96 |
| C381 | 0.95 |
| C120 | 1 |
| C781 | 0.86 |
| C612 | 0.94 |
| C772 | 0.93 |
| C042 | 0.99 |
| C017 | 0.97 |

**Table S3.**
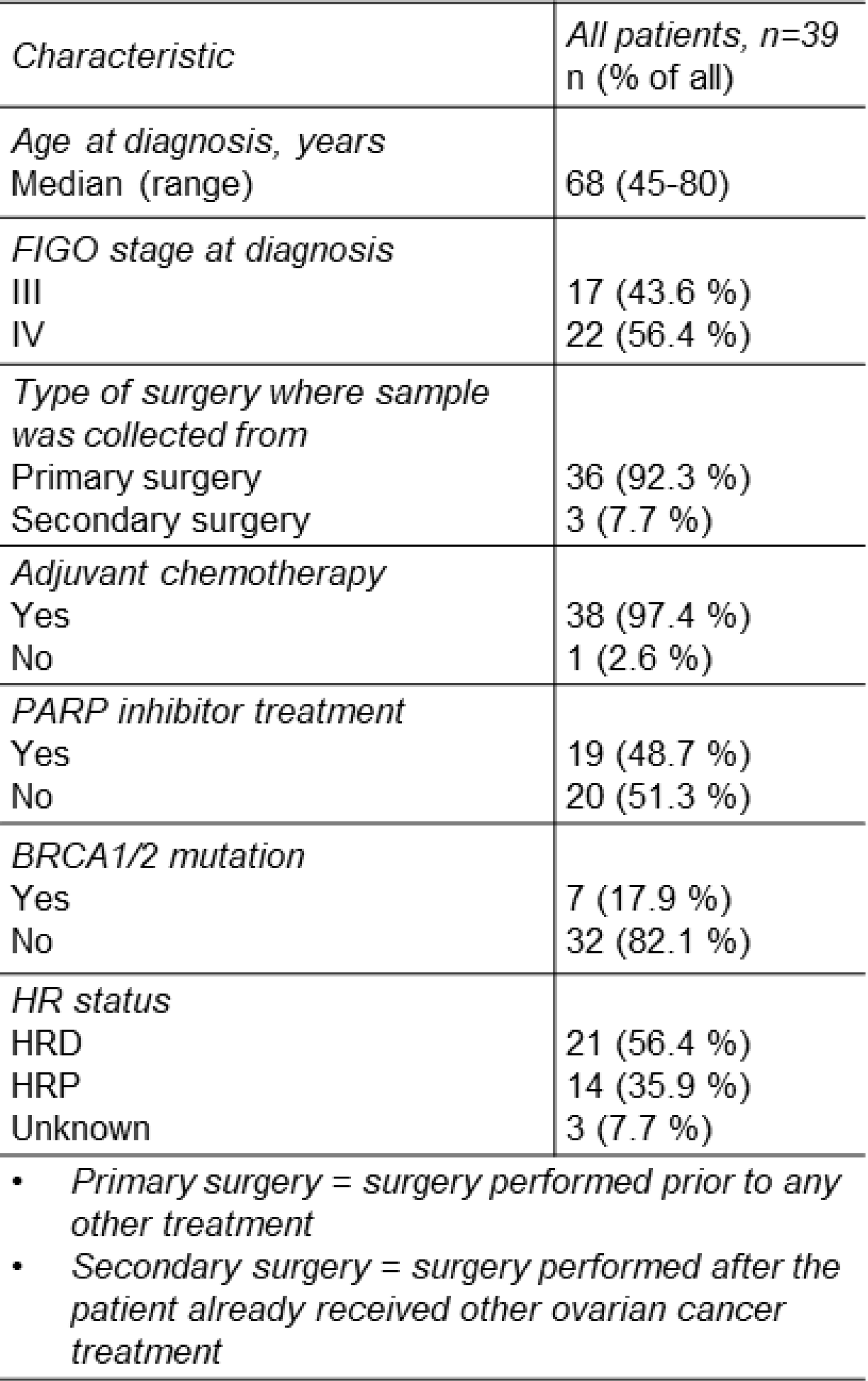
Summary of clinical characteristics of patients included in this study.

| *Characteristic* | *All patients, n=39*<br>*n (% of all)* |
| --- | --- |
| *Age at diagnosis, years*<br>Median (range) | 68 (45-80) |
| *FIGO stage at diagnosis*<br>III<br>IV | 17 (43.6 %)<br>22 (56.4 %) |
| *Type of surgery where sample*<br>*was collected from*<br>Primary surgery<br>Secondary surgery | 36 (92.3 %)<br>3 (7.7 %) |
| *Adjuvant chemotherapy*<br>Yes<br>No | 38 (97.4 %)<br>1 (2.6 %) |
| *PARP inhibitor treatment*<br>Yes<br>No | 19 (48.7 %)<br>20 (51.3 %) |
| *BRCA1/2 mutation*<br>Yes<br>No | 7 (17.9 %)<br>32 (82.1 %) |
| *HR status*<br>HRD<br>HRP<br>Unknown | 21 (56.4 %)<br>14 (35.9 %)<br>3 (7.7 %) |
| *Primary surgery = surgery performed prior to any other treatment*<br>*Secondary surgery = surgery performed after the patient already received other ovarian cancer treatment* |  |

**Table S4.** Details of compounds tested for single and combinatorial treatments.

| TARGET | COMPOUND | CATALOG # | COMPANY | SOLVENT | STOCK<br>CONCENTRATION (mM) | SINGLE AGENTS TESTED (µM) |
| --- | --- | --- | --- | --- | --- | --- |
| PARP | Olaparib | S1060 | Selleckchem | DMSO | 10mM | 10 |
|  |  |  |  |  |  | 23 |
|  |  |  |  |  |  | 37 |
|  |  |  |  |  |  | 50 |
| ATR | Berzosertib (VE-822) | S7102 | Selleckchem | DMSO | 10 mM | 1 |
|  |  |  |  |  |  | 4 |
|  |  |  |  |  |  | 7 |
|  |  |  |  |  |  | 10 |
| WEE1 | Adavoseritb | S1525 | Selleckchem | DMSO | 10 mM | 1 |
|  |  |  |  |  |  | 4 |
|  |  |  |  |  |  | 7 |
|  |  |  |  |  |  | 10 |
| Autotaxin (ATX) | Ziritaxestat (GLPG1690) | HY-101772 | MedChemExpress | DMSO | 1mM | 0.2 |
|  |  |  |  |  |  | 0.5 |
|  |  |  |  |  |  | 0.7 |
|  |  |  |  |  |  | 1 |
| PERK | AMG PERK 44 | HY-12661A | MedChemExpress | DMSO | 1mM | 0.2 |
|  |  |  |  |  |  | 0.5 |
|  |  |  |  |  |  | 0.7 |
|  |  |  |  |  |  | 1 |
| PD-1 | Pembrolizumab | A2005 | Selleckchem | PBS | 5 mg/mL | 10 mg/mL |
|  |  |  |  |  |  | 23 mg/mL |
|  |  |  |  |  |  | 37 mg/mL |
|  |  |  |  |  |  | 50 mg/mL |

| Combination | COMPOUND | Concentration |
| --- | --- | --- |
| 1 | Olaparib | 10 µM |
|  | Pemnnrolizumab | 10 µg/ml |
| 2 | Olaparib | 36 µM |
|  | Pemnnrolizumab | 36 µg/ml |
| 3 | Olaparib | 10 µM |
|  | Ziritaxestat (GLPG1690) | 0.2 µm |
| 4 | Olaparib | 36 µM |
|  | Ziritaxestat (GLPG1690) | 0.7 µm |
| 5 | Olaparib | 10 µM |
|  | AMG PERK 44 | 0.2 µm |
| 6 | Olaparib | 36 µM |
|  | AMG PERK 44 | 0.7 µm |
| 7 | Olaparib | 10 µM |
|  | Berzosertib (VE-822) | 1 µM |
| 8 | Olaparib | 36 µM |
|  | Berzosertib (VE-822) | 7 µM |
| 9 | Olaparib | 10 µM |
|  | Adavoseritb | 1 µm |
| 10 | Olaparib | 36 µM |
|  | Adavoseritb | 7 µm |
| 11 | Berzosertib (VE-822) | 1 µM |
|  | Ziritaxestat (GLPG1690) | 0.2 µm |
| 12 | Berzosertib (VE-822) | 10 µM |
|  | Ziritaxestat (GLPG1690) | 0.7 µm |
| 13 | Berzosertib (VE-822) | 1 µM |
|  | AMG PERK 44 | 0.2 µm |
| 14 | Berzosertib (VE-822) | 10 µM |
|  | AMG PERK 44 | 0.7 µm |
| 15 | Berzosertib (VE-822) | 1 µM |
|  | Adavoseritb | 1 µm |
| 16 | Berzosertib (VE-822) | 7 µM |
|  | Adavoseritb | 7 µm |
| 17 | Adavoseritb | 1 µm |
|  | Ziritaxestat (GLPG1690) | 0.2 µm |
| 18 | Adavoseritb | 7 µm |
|  | Ziritaxestat (GLPG1690) | 0.7 µm |
| 19 | Adavoseritb | 1 µm |
|  | AMG PERK 44 | 0.2 µm |
| 20 | Adavoseritb | 7 µm |
|  | AMG PERK 44 | 0.7 µm |
| 21 | Adavoseritb | 1 µm |
|  | Pembrolizumab | 10 µg/ml |
| 22 | Adavoseritb | 7 µm |
|  | Pembrolizumab | 36 µg/ml |
| 23 | Pembrolizumab | 10 µg/ml |
|  | Ziritaxestat (GLPG1690) | 0.2 µm |
| 24 | Pembrolizumab | 36 µg/ml |
|  | Ziritaxestat (GLPG1690) | 0.7 µm |
| 25 | Pembrolizumab | 10 µg/ml |
|  | AMG PERK 44 | 0.2 µm |
| 26 | Pembrolizumab | 36 µg/ml |
|  | AMG PERK 44 | 0.7 µm |
| 27 | Pembrolizumab | 10 µg/ml |
|  | AMG PERK 44 | 0.2 µm |
| 28 | Pembrolizumab | 36 µg/ml |
|  | AMG PERK 44 | 0.7 µm |
| 29 | Ziritaxestat (GLPG1690) | 0.2 µm |
|  | AMG PERK 44 | 0.2 µm |
| 30 | Ziritaxestat (GLPG1690) | 0.7 µm |
|  | AMG PERK 44 | 0.7 µm |

**Fig. S1.**
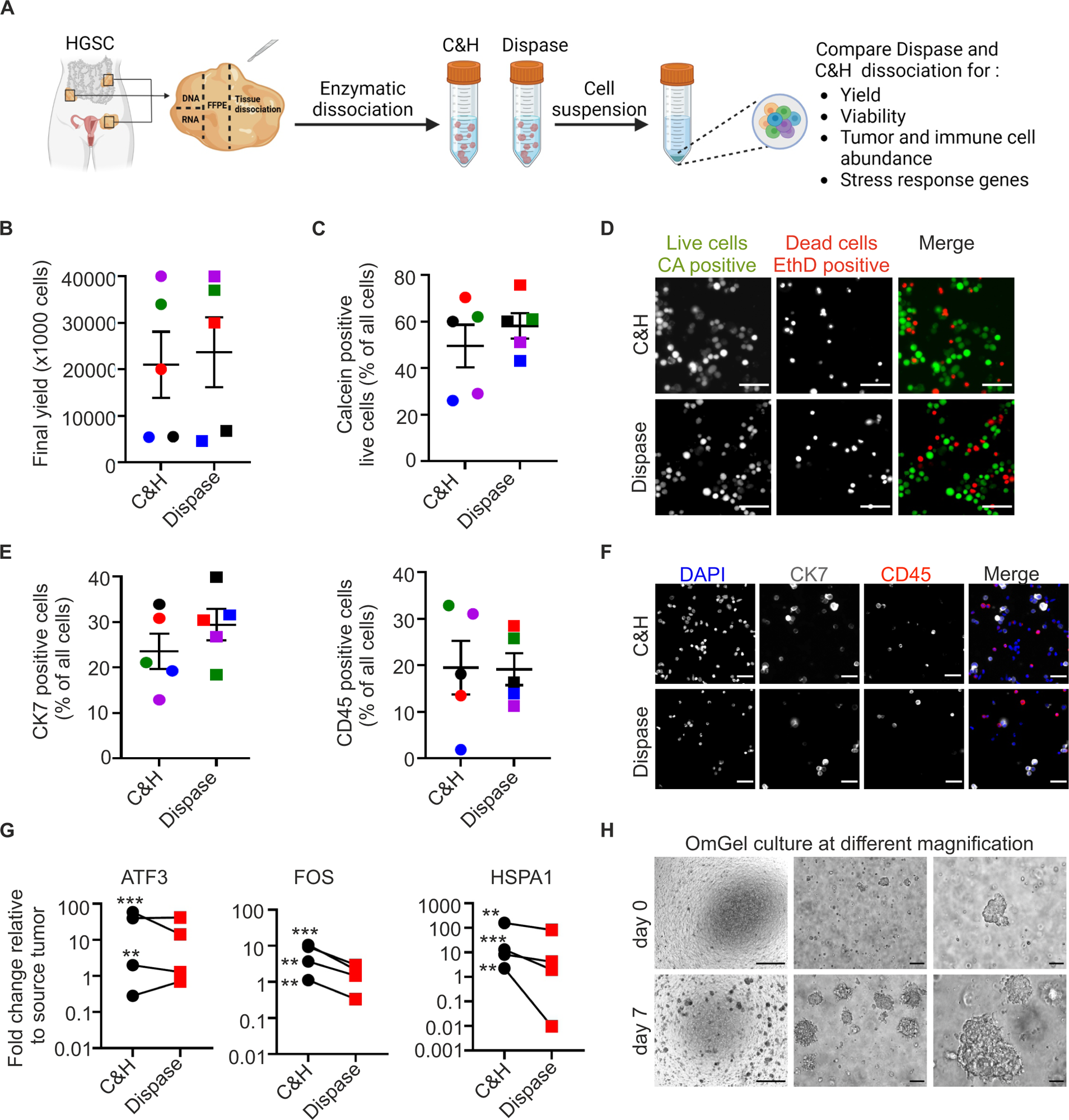
Nature of the enzymatic tissue dissociation influence on the expression of stress response genes. **A).** Workflow for evaluating the effect of enzymatic dissociation on the final yield, recovery of tumor and immune cells, and expression of stress response genes. **B) & C).** Quantification of the final yield, and cell viability following C&H or dispase dissociation, each color represents an individual sample. **D).** Representative IF images showing calcein (CA) positive live cells or Ethidium homodimer-1 (EthD) positive dead cells from C&H or dispase dissociation. Scale bar 50µM. **E).** Quantification of % of CK7 or CD45 positive cells following C&H or dispase dissociation, each color represents an individual sample. **F).** Representative IF images of the cells dissociated with C&H or dispase and stained with CK7 or CD45 antibodies. Scale bar 50µM. **G)**. qPCR analysis showing the expression of stress response genes *ATF3*, *FOS*, and *HSPA1* in the cells dissociated with C&H or dispase. **H).** Representative brightfield images of d0 and d7 OmGel cultures at different magnifications. Scale bar from left to right: 1mm, 100µM, 50µM.

**Fig. S2.**
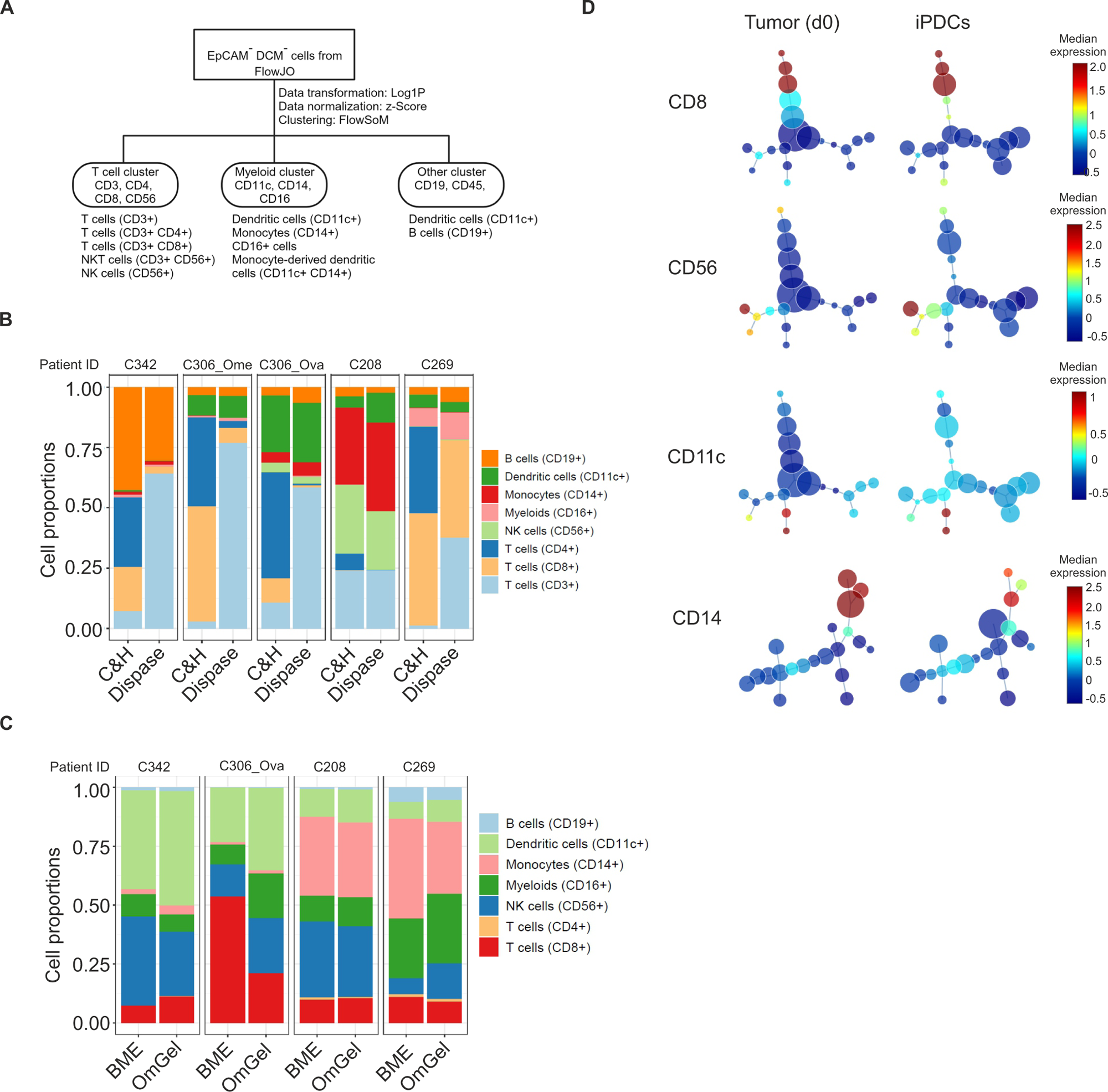
HGSC iPDCs maintain source tumor-derived immune cells and show HRD genotype-specific functional immune cell responses to Olaparib treatment. **A).** Flowchart describing the steps involved in flow cytometry data using CYTO. EpCAM^-^ live cells gated from the FlowJo are analyzed in a hierarchical manner to finally annotated into different cell types. **B).** Bar plot showing the proportions of indicated immune cells following C&H and dispase dissociation. **C).** Bar plot showing the proportions of indicated immune cells in iPDCs cultured using BME and OmGel for 4-6 days. **D).** Minimum spanning tree showing median marker expression in tumor and matched iPDCs. Each dot represents individual clusters, color and size of the dots indicates median marker expression, and cluster size, respectively.

**Fig. S3.**
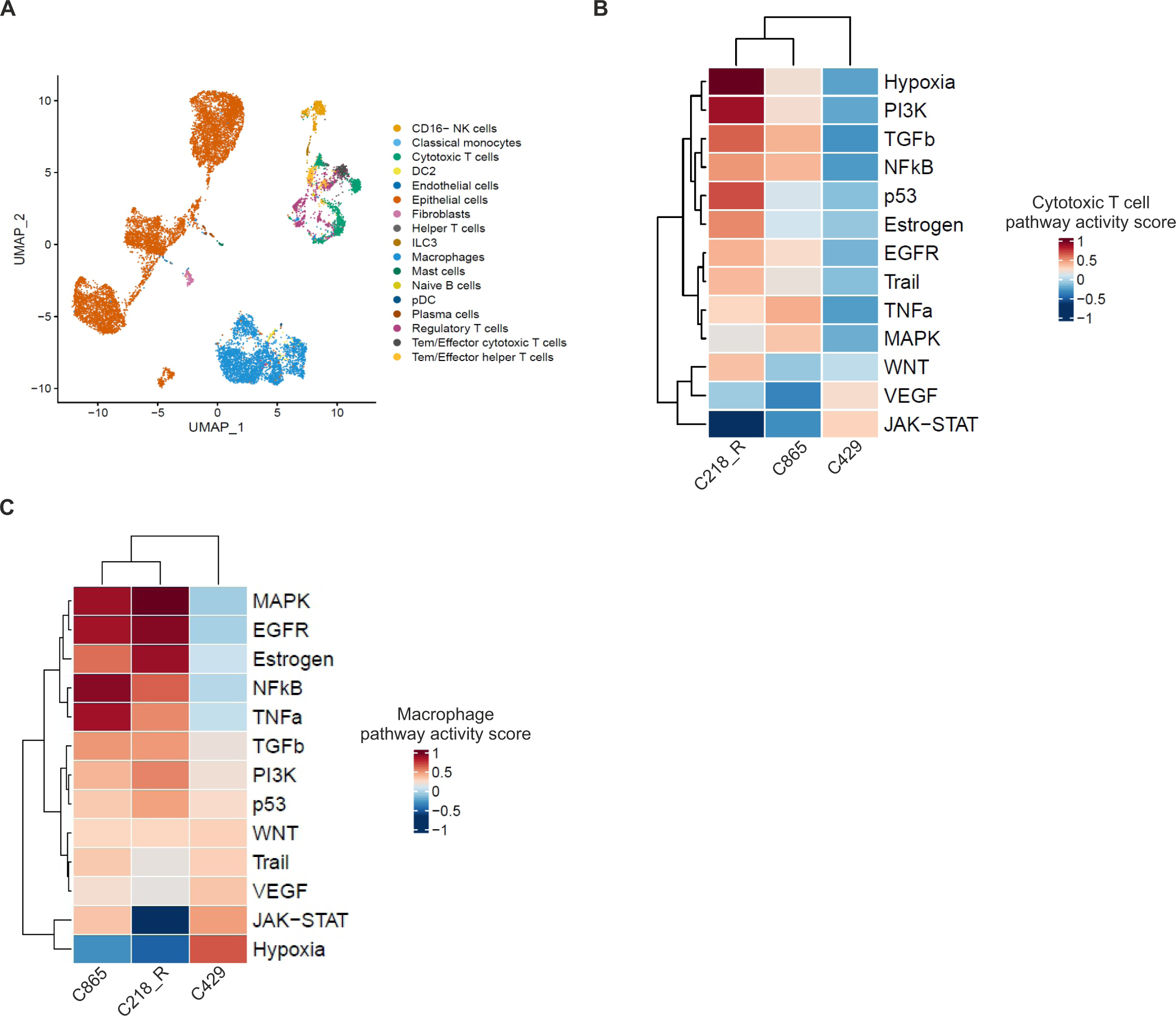
PARPi and chemotherapy resistant tumors show distinct pathway activity across different immune cell types. **A).** UMAP plot showing epithelial and immune cell type clusters from the tumors of three resistant patients analyzed by scRNAseq. **B).** Heatmap showing the pathway activity scores in CD8+ T cells and macrophages **C).** from the tumors of resistant patients.

**Fig. S4.**
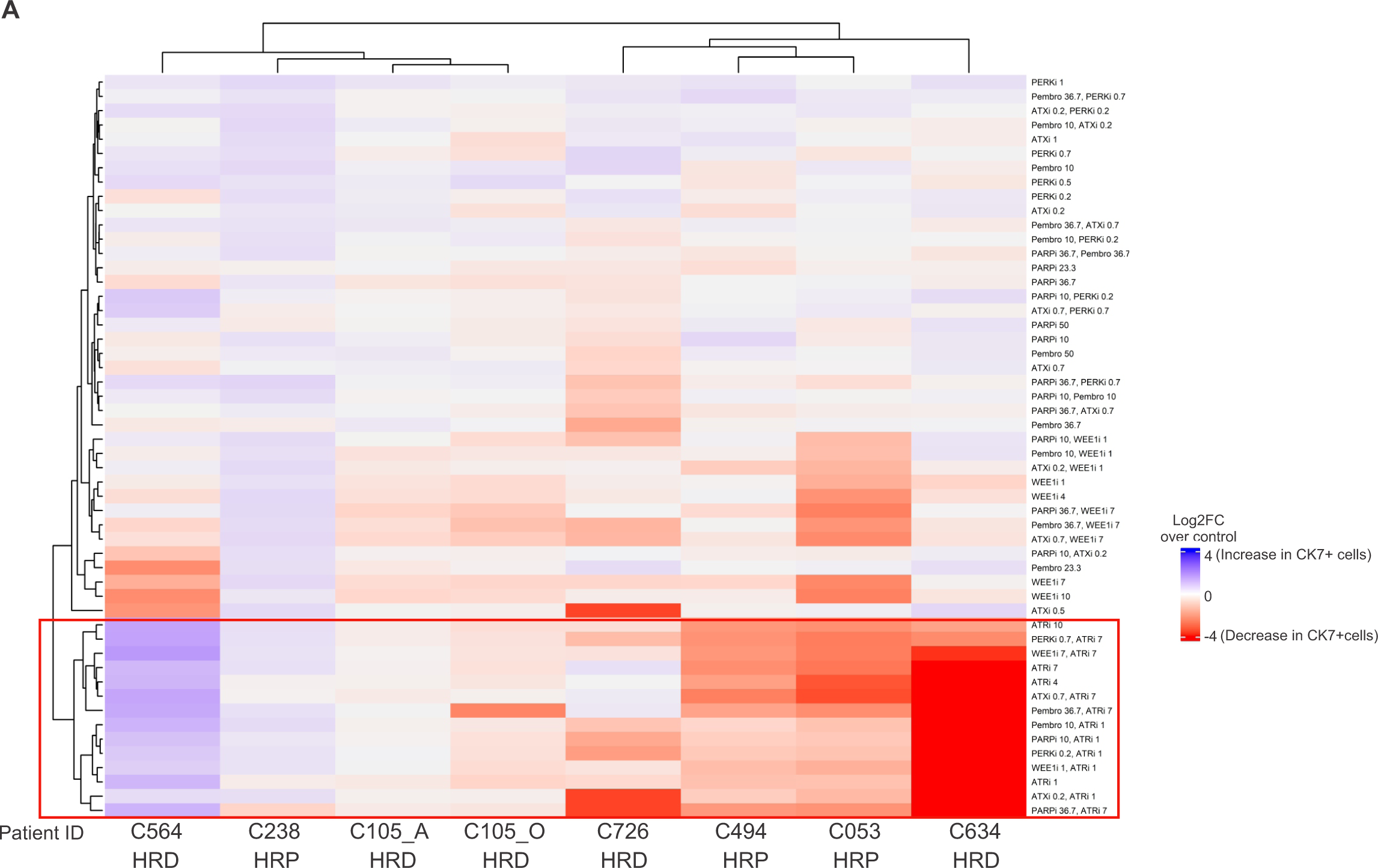
iPDCs provide a platform for high-throughput drug testing. **A). Heatmap showing the** unsupervised hierarchical clustering of Log2FC in live CK7+ tumor cells over all live cells in each treatment condition normalized to control, across 8 different tumor-derived iPDCs.

